# Homeostasis of the intervertebral disc requires regulation of STAT3 signaling by the adhesion G-protein coupled receptor ADGRG6

**DOI:** 10.1101/581595

**Authors:** Zhaoyang Liu, Garrett W.D. Easson, Jingjing Zhao, Nadja Makki, Nadav Ahituv, Matthew J. Hilton, Simon Y. Tang, Ryan S. Gray

## Abstract

Degenerative changes of the intervertebral disc (IVD) are a leading cause of disability affecting humans worldwide. While this is primarily attributed to trauma and aging, genetic variation is associated with disc degeneration in humans. However, the precise mechanisms driving the initiation and progression of disease remain elusive due to a paucity of genetic animal models. Here, we discuss a novel genetic mouse model of endplate-oriented disc degeneration. We show that the adhesion G-protein coupled receptor G6 (ADGRG6) mediates several anabolic and catabolic factors, fibrotic collagen genes, pro-inflammatory pathways, and mechanical properties of the IVD, prior to the onset of overt histopathology of these tissues. Furthermore, we found increased IL-6/STAT3 activation in the IVD and demonstrate that treatment with a chemical inhibitor of STAT3 activation ameliorates disc degeneration in these mutant mice. These findings establish ADGRG6 as a critical regulator of homeostasis of adult disc homeostasis and implicate ADGRG6 and STAT3 as promising therapeutic targets for degenerative joint diseases.

**Author summary:** Degenerative changes of the intervertebral disc (IVD) are a leading cause of disability affecting humans worldwide. While this is primarily attributed to trauma and aging, genetic variation is associated with disc degeneration in humans. However, the precise mechanisms driving the initiation and progression of disease remain elusive due to a paucity of genetic animal models. Here, we discuss a novel genetic mouse model of endplate-oriented disc degeneration. We show that the adhesion G-protein coupled receptor G6 (ADGRG6) mediates fibrotic collagen expression, causing increased mechanical stiffness of the IVD prior to the onset of histopathology in adult mice. Furthermore, we found increased IL-6/STAT3 activation in the IVD and demonstrate that treatment with a chemical inhibitor of STAT3 activation ameliorates disc degeneration in these mutant mice. Our results demonstrate that ADGRG6 regulation of STAT3 signaling is important for IVD homeostasis, indicating potential therapeutic targets for degenerative joint disorders.

## Introduction

Spine disorders are one of the most common health issues affecting human populations worldwide, causing a tremendous socio-economic burden. The progression of spine disorders such as low back pain, disc herniation, endplate fracture, and scoliosis are associated with degenerative changes of the intervertebral disc (IVD) (Brinjikji et al., 2015; Dudli et al., 2014; Sun et al., 2017; Zhu et al., 2017b). Therefore, elucidation of the pathways and signaling important for maintaining spine stability and the development and homeostasis of the IVD tissues is critical for the diagnosis, prevention, and treatment of degenerative spine disorders.

The IVD is a fibrocartilaginous joint that connects two adjacent vertebrae and provides structural stability, flexibility, and cushions axial loading of the spinal column (Cortes and Elliott, 2014). The disc is composed of a proteoglycan-rich nucleus pulposus, surrounded by a multi-lamellar annulus fibrosus, and situated between the cartilaginous endplates, which provide nutritional flux to the IVD (Figure 1A). Hallmarks of disc degeneration in humans include loss of disc height, reduced proteoglycan staining, and accumulation of markers of fibrosis within the disc. At the same time, the cartilaginous endplate may show signs of degeneration and calcification, which further compromises nutrient availability to the inner disc layers (Vergroesen et al., 2015). More severe forms of disc degeneration can also result in the herniation of the nucleus pulposus (i) laterally through the annulus fibrosis layer; or (ii) through the cartilaginous endplate into the vertebral body (endplate-oriented), clinically referred to as Schmorl’s nodes (Fardon et al., 2014). Genetic susceptibility to disc degeneration has been shown to play a major role in disc degeneration (Sambrook et al., 1999), with the majority of these findings implicating extracellular matrix components of the disc, matrix metalloproteases, or pro-inflammatory cytokines (Munir et al., 2018). Together these data suggest that dysregulation of anabolic and catabolic factors as well as inflammatory signaling may underlie many forms of disc degeneration in humans. However, the molecular regulators and initiating factors for these events remain to be defined.

**Figure 1:**
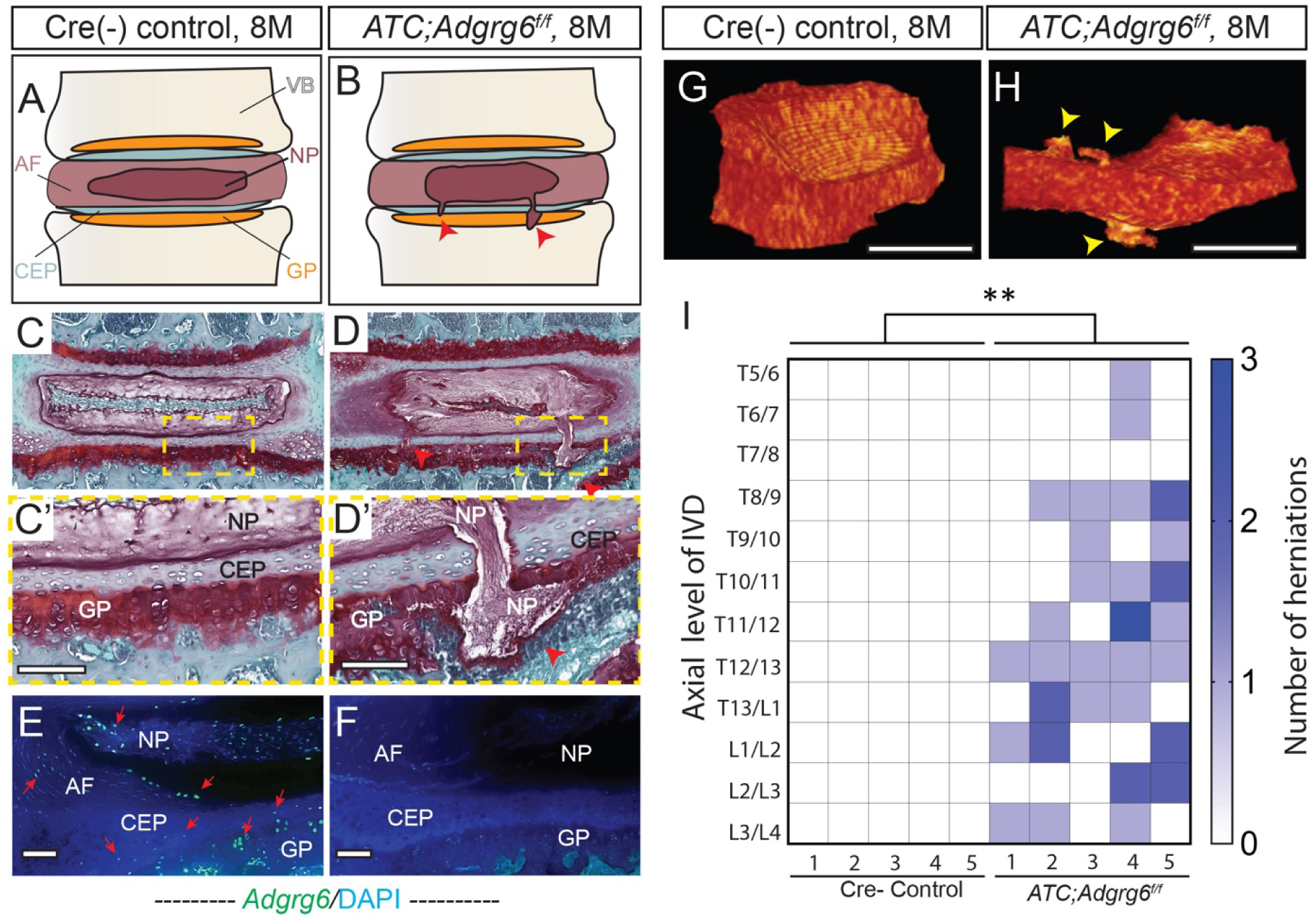
Adult *ATC;Adgrg6^f/f^* mutant mice display endplate-oriented herniations of the IVD. (A and B) Schematic of endplate-oriented herniations (B, red arrowheads) observed in *ATC;Adgrg6^f/f^* mutant mice (B), in contrast to a typical wild-type IVD (A) at 8 months of age. (C-D’) Representative medial-sectioned 8-month-old mouse IVDs stained with Safranin-O/Fast green (SO/FG) (*n*=3 for each group). (E, F) *Adgrg6* riboprobe FISH (green fluorescence) at 8 months. (G, H) Representative reconstructions of contrast-enhanced μCT of Cre (-) control (G) and *ATC;Adgrg^f/f^* mutant (H) IVDs at 8 months of age. Endplate-oriented herniations are observed in SO/FG stained sections (D’, red arrowheads) and by contrast-enhanced μCT (H, yellow arrowheads) (*n*=5 for each group). (I) Heat map of contrast-enhanced μCT data from five Cre (-) control and five *ATC;Adgrg^f/f^* mutant mouse spines, plotting the axial level of the IVD (left axis) and the number of herniations (right axis) observed in each mouse. (**p≤0.01, Student’s *t* Test.) Scale bars: 50μm in (C’, D’), 100μm in (E, F), and 500μm in (G, H). *AF-annulus fibrosis, CEP-cartilaginous endplate, GP-growth plate, NP-nucleus pulposus, and VB-vertebral body*.

Here, we show that ADGRG6 has a critical role in intervertebral disc homeostasis through regulating STAT3 signaling. ADGRG6 (also called GPR126) is a member of the adhesion G-protein coupled receptor (aGPCR) family of proteins, all of which are thought to have a canonical intercellular signaling function via G-protein coupled signaling, as well as a potential for cell-cell or cell-matrix signaling via the extracellular N-terminal fragment (Patra et al., 2013). In zebrafish, *adgrg6/gpr126* is critical for the development of cartilaginous tissues of the semicircular canal via regulation extracellular matrix (ECM) gene expression (Geng et al., 2013), suggesting a role for ADGRG6 in the regulation of cartilaginous tissues. Conditional loss of *Adgrg6* in multipotent osteochondral progenitors-giving rise to bone, cartilage, and some connective tissues- of the spine generated postnatal-onset scoliosis, ribcage deformity, and led to midline clefts in the endplates and annulus fibrosus (Karner et al., 2015). Since the development of scoliosis is often linked with IVD deformity (Zhu et al., 2017a), we sought to investigate the role of ADGRG6 in specifically in cartilaginous tissues of the IVD during embryonic and postnatal development.

To define the role of ADGRG6 we combined conditional mouse genetics, genomic approaches, mechanical assessment of intervertebral disc, and cell biological approaches in chondrogenic cell culture. Together, these studies reveal that ADGRG6 has a conserved role for the maintenance of normal chondrogenic gene expression profiles and regulation of STAT3 signaling. We demonstrate that loss of ADGRG6 leads to increased expression of fibrotic collagens and alteration of the normal biomechanical properties of the IVD, prior to the onset of endplate-oriented herniations. Finally, we demonstrate loss of ADGRG6 leads to increased, ectopic STAT3 activation in the IVD and that blockade of STAT3 activation can alleviate some degenerative changes of the IVD and progression of endplate-oriented IVD herniation in *Adgrg6* conditional mutant mice. Taken together, our work establishes ADGRG6 as a novel regulator of IVD endplate integrity in the mouse and suggests that modulation of ADGRG6/STAT3 signaling could provide robust disease-modifying targets for endplate-oriented disc degeneration, in humans.

## Results

### Loss of ADGRG6 in intervertebral discs leads to endplate-oriented herniations in adult mice

We have found that conditional removal of ADGRG6 function in the intervertebral disc (IVD) results in the initiation and progression specifically of cartilaginous endplate defects in mice (Figure 1B). *Adgrg6* is robustly expressed in the growth plate, but not in cortical or trabecular bone in vertebrae of adult mice by *in situ* hybridization (Supplemental Figure 1A). The more sensitive fluorescent *in situ* hybridization (FISH) detection method also established *Adgrg6* expression in cells of the cartilaginous endplate, annulus fibrosus, and nucleus pulposus (Supplemental Figure 1C and Figure 1E). To determine the role of ADGRG6 function specifically in these committed chondrogenic lineages of the spine, we utilize an <u>A</u>ggrecan enhancer-driven, <u>T</u>etracycline-inducible <u>C</u>re (*ATC*) transgenic mouse strain (Dy et al., 2012) (*ATC;Adgrg6^f/f^*). We established two experimental groups to address the temporal requirement of ADGRG6 function by induction during embryonic development (from E0.5-P20, prior to IVD specification) or during perinatal development (from P1-P20, after IVD specification).

For both induction strategies, we consistently observed endplate-oriented disc herniations (Adams et al., 2012) in adult mutant mice (Figure 1 and Supplemental Figure 2). *ATC;Adgrg6^f/f^* mutant mice induced during perinatal developmental displayed prolapse of the nucleus pulposus material into the vertebra body (Figure 1B, D, D’), as did mutant mice recombined during embryonic development (Supplemental Figure 2B, B’) (*n=*3 mice for perinatal induction group; *n*=6 mice for embryonic induction group). These data, suggest that the role of ADGRG for the initiation and progression of IVD defects can largely be attributed to its function during postnatal development. Interestingly, histological analysis of the IVD of *ATC;Adgrg6^f/f^* mutant mice after either induction strategy did not reveal obvious changes in disc height, alterations of overall sulfated-proteoglycan abundance (Safranin-O staining) (Figure 1D, Supplemental Figure 2), or defects of the annulus fibrosis or nucleus pulposus tissues, suggesting a unique endplate-driven pathology for this model.

We found that traditional two-dimensional histological analysis limited our ability to capture the extent and distribution of these endplate-oriented herniations along the spine. To address this, we exploited contrast-enhanced micro-computed tomography (μCT) imaging which allows for a full three-dimensional analysis and segmented visualization of the IVD within the intact mouse spine (Lin and Tang, 2017). Reconstruction and segmentation of the whole IVD in 8-month-old *ATC;Adgrg6^f/f^* mutant (Figure 1H; Movie 1) and Cre (-) control mice (Figure 1G; Movie 2) (49 discs total; *n*=5 mice/genotype) revealed multiple incidences of endplate-oriented disc herniations (yellow arrowheads, Figure 1H). Quantification of the contrast-enhanced μCT imaging indicated from 0-3 herniations present/IVD in *ATC;Adgrg^f/f^* mutant mice, while Cre (-) control mice did not demonstrate similar defects (Figure 1I). Spatial analysis shows that these herniations occurred along the entire axis of the mutant spine that was imaged (Thoracic (T)5/6 – Lumbar (L)3/4), without obvious hotspots. As above, we did not observe radial fissures or lateral prolapse of the disc in *ATC;Adgrg6^f/f^* mutant mice using this imaging approach, which further supports that IVD degeneration in these conditional mutant mice are specifically occurring by endplate-driven mechanisms (Adams et al., 2012).

Our previous work demonstrated a clear role for *Adgrg6* in the formation of late-onset scoliosis in mouse (Karner et al., 2015), however the cellular pathogenesis of this process remains unresolved. Interestingly, while ∼80% of *Col2Cre;Adgrg6^f/f^* mutant mice demonstrated late-onset scoliosis (Karner et al., 2015), we only observed scoliosis in ∼12% of *ATC;Adgrg6^f/f^* mutant mice (not shown). One possible explanation is the difference in targeted tissues between the two Cre deleter strains. Analysis of recombination using β-galactosidase staining of *ATC;Rosa-LacZ* mice showed nearly complete recombination throughout the IVD and growth plate at weaning regardless of the timing of induction (Supplemental Figure 3C, D). However, the outer most layers of the annulus fibrosus and the periosteum were not targeted using either strategy (Supplemental Figure 3A, C, D). In contrast, the entire IVD as well as periosteum and trabecular bone in the vertebral body is completely recombined at P1 when crossed to the *Col2Cre* deleter strain, with marked reduction of recombination in these tissues using the *ATC* deleter strain (Supplemental Figure 3B, E). However, effective knockdown of *Adgrg6* in *ATC;Adgrg6^f/f^* mutant mice was further confirmed by FISH analysis at 8 months (Figure 1F, Supplemental Figure 1B, D) and by qPCR analysis of gene expression in IVDs extracted from 1.5 month-old mice (Figure 2O). *In situ* hybridization analysis showed that *Adgrg6* expression in non-targeted tissues such as the periosteum was not altered (Supplemental Figure 1E, F). These data suggest that the IVD is important for susceptibility of scoliosis, yet additional effectors of spine stability are involved. However, *ATC;Adgrg^f/f^* mutant mice consistently exhibited endplate-oriented herniations irrespective of whether thoracic scoliosis was observed (Figure 1I and Liu and Gray, unpublished data). These data demonstrate that ADGRG6 has a unique role in the regulation of postnatal homeostasis of the IVD, in addition to scoliosis.

**Figure 2:**
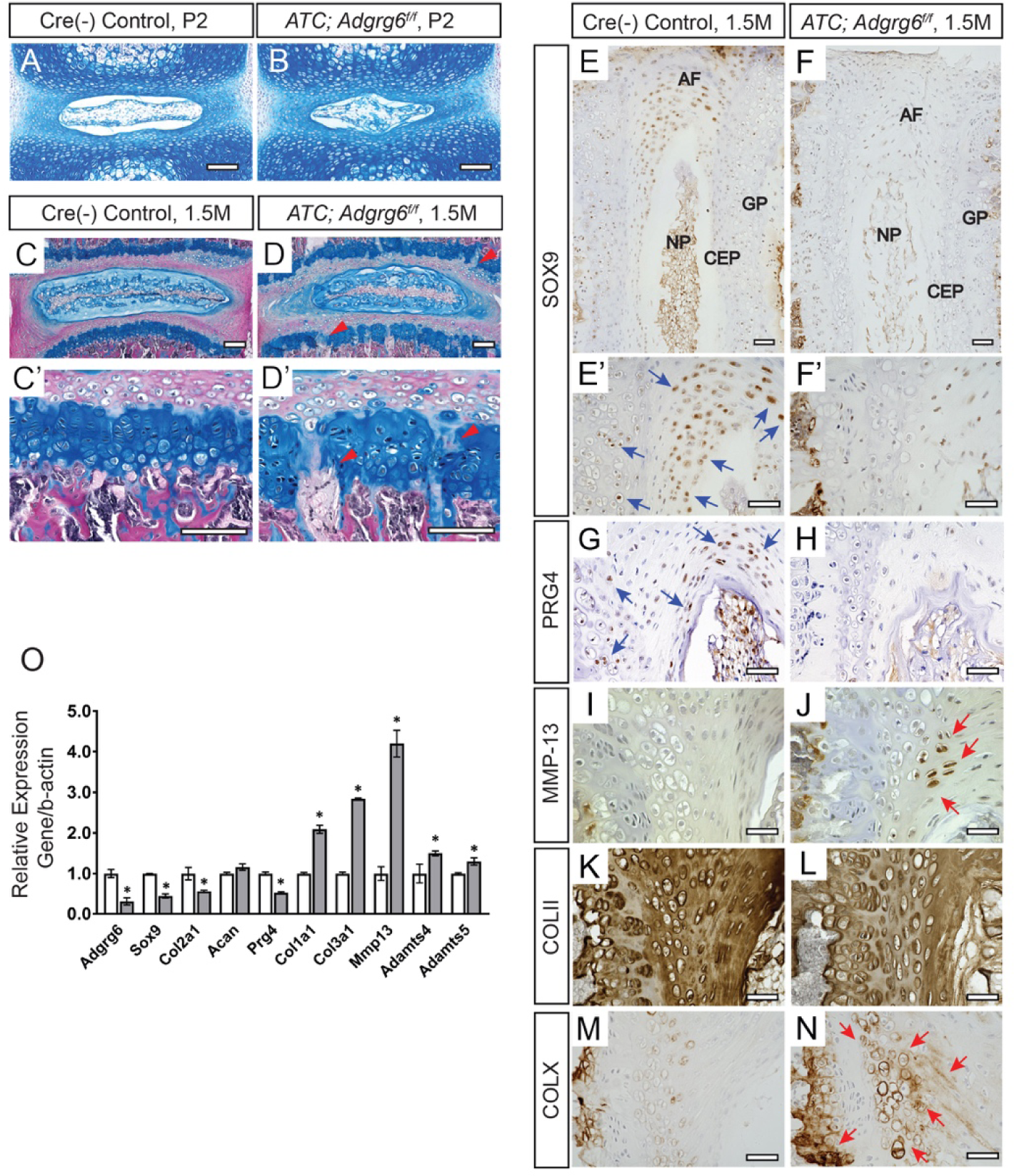
Young *ATC;Adgrg6^f/f^* mutant mice display alterations in IVDs consistent with disc degeneration pathology. (A-B’) Representative medial-sectioned IVD tissues stained with Alcian blue-Hematoxylin/Orange G. *ATC;Adgrg^f/f^* mutant and control IVDs are comparable at P2 (A, B) and 1.5 months (C-D’) of age, except for the mild increase of acellular clefts in the GP at 1.5 months (red arrowheads, D, D’ *n*=3 for each group.) (E-O) IHC (E-N) and qPCR (O) analyses of common markers of degenerative disc in mice at 1.5 months. *ATC;Adgrg^f/f^* mutant IVDs display reduced expression of markers of healthy disc: SOX9/*Sox9* (blue arrows, E’) and PRG4/*Prg4* (blue arrows, G), and mildly reduced COLII/*Col2a1* (K). They also display increased expression of hypertrophic marker COLX/*Col10a1* (red arrows, N), extracellular matrix modifying enzymes (MMP-13/*Mmp13* (red arrows, J), *Adamts4*, and *Adamts5*), de-differentiation marker (*Col1a1*), and fibrosis marker (*Col3a1*). (C-N, n= 3 for each group. O, *n*= 3 biological replicates. Bars represent mean ± SD. *p≤0.05, Student’s *t* Test.) Scale bars: 100μm in (A, B) and (C’, D’); 200μm in (C, D); 50μm in (E-N). *AF-annulus fibrosis, CEP-cartilaginous endplate, GP-growth plate, and NP-nucleus pulposus*.

### Loss of ADGRG6 in the intervertebral discs leads to alterations of the extracellular matrix, consistent with human disc degeneration pathology

We next sought to understand the molecular mechanism underlying the initiation of discogenic defects. To guide our analysis, we took cues from molecular changes observed in degenerative human IVDs (Yee and Chan, 2014), and revealed several indicators of degenerative joint disease in *ATC;Adgrg6^f/f^* mutant mice at 1.5 months, prior to overt histopathology (Figure 2D, D’, induced from E0.5-P20). Using immunohistochemistry (IHC) we observed reduction in the expression of the transcription factor SOX-9 (SOX9) (Figure 2F, F’ and Supplemental Figure 5B) and proteoglycan 4 (PRG4) (Figure 2H and Supplemental Figure 5D) in the inner layer of annulus fibrosus and endplate of the mutant mice. At the same time, expression of the hypertrophic chondrocyte marker type X collagen (COLX) was increased in the endplate and growth plate of the mutant IVD (Figure 2N and Supplemental Figure 5J). We also observed a minor reduction of type II collagen (COLII) staining (Figure 2L and Supplemental Figure 5H) and increased, ectopic expression of matrix metalloprotease-13 (MMP-13) protein in the mutant mice (Figure 2J and Supplemental Figure 5F). Analysis of *ATC;Adgrg6^f/f^* mutant mice induced after birth (P1-P20) demonstrated similar alterations in protein expression in the IVD (Supplemental Figure 4C-J). These results further support the postnatal role for ADGRG6 function in degenerative process of the IVD. Alterations of protein expression were supported by changes in gene expression (Figure 2O), where we observed reduced expression of *Col2a1*, *Sox9* and *Prg4* expression in *ATC;Adgrg^f/f^* mutant mice at 1.5 months. Interestingly, the expression of several catabolic mediators including *Mmp13*, *Adamts4*, and *Adamts5,* as well as markers of de-differentiation and fibrosis, *Col1a1* and *Col3a1,* were increased in the mutant mice. Together, these alterations of typical gene expression profiles in the IVD are consistent with expression profiles reported for degenerative joint diseases, including degenerative discs (Kadow et al., 2015) and osteoarthritis of articulating joints (Kalb et al., 2012; Tonge et al., 2014) in humans.

As discussed above, ADGRG6 is seemingly not critical for early pattering and morphology of cartilaginous tissues of IVD, rather it appears to regulate postnatal and adult developmental processes of these tissues. In agreement, we show that embryonic loss of *Adgrg6* in cartilaginous tissues does not lead to obvious alternations of the Alcian blue staining or dysmorphology of the disc at P2 (Figure 2B) or at 1.5 months of age (Figure 2D). However, we did observe a mild increase in appearance of acellular clefts in the mutant growth plate (7.2±3.4 clefts/IVD; *n*=4; *p*=0.03) (Figure 2D-D’) at 1.5 months, compared to littermate controls (2.5±0.4 clefts/IVD; *n*=4) (Figure 2C-C’). This is consistent with our observations of *ATC;Adgrg6^f/f^* mutant mice, recombined postnatally, which demonstrate minor defects in growth plate at 4-months (40%; *n*=5) (yellow arrows, Supplemental Figure 4B). In addition, TUNEL staining demonstrated a mild increase in cell death in *ATC;Adgrg6^f/f^* mutant mice within the annulus fibrosus, nucleus pulposus, and endplate compartments of the IVD, less so within vertebral growth plate at 1.5 months (Supplemental Figure 6B, C). Taken together these data strongly support the critical role of ADGRG6 function in cartilaginous tissues of the IVD during postnatal/adult developmental processes.

### Mechanical properties are altered prior to the onset of obvious histopathology in the intervertebral discs of ADGRG6 deficient mice

During the progression of disc degeneration and osteoarthritis-related joint remodeling in humans, increased catabolic factors, inflammation, coupled with alterations of normal extracellular matrix composition of the IVD can have deleterious effects on the mechanical properties on these tissues (Nguyen et al., 2017). In order to assess alterations of mechanical properties of *ATC;Adgrg6^f/f^* mutant IVDs, we isolated individual lumbar discs (Lum.1/2 and 4/5) for dynamic mechanical testing (16 discs total; *n*=4 mice/genotype) from 1.5-month-old mutant and control mice. Using micro-indentation we demonstrated a consistent increase in the stiffness (Newton (N)/mm) of *ATC;Adgrg6^f/f^* mutant IVDs under 5% strain cyclic loading (Figure 3A; *p*=0.0114; mean w/95% CI) and under 50% monotonic overloading (Figure 3B; *p*=0.0026; mean w/95% CI). Stiffening of the IVD is commonly observed with early-onset degenerative changes, which compromises the damage resistance of the tissue (Liu et al., 2015a). We analyzed proteoglycan quantification in these IVDs by dimethylmethylene blue assay and found no significant alterations comparing mutant and littermate control mice at 1.5 months. However, we did observed a mild increase in total collagen content in *ATC;Adgrg6^f/f^* mutant IVDs by hydroxyproline assay (measured as collagen/wet weight; *p* = 0.0561; one-tailed *t*-test), which is consistent with the elevated levels of multiple fibrotic collagen expression in 1.5-month-old mutant IVDs (Figure 2O). We speculate that, alterations in normal extracellular matrix/collagen gene and protein expression, coupled with increased cell death compromise a constellation of factors leading to the decline in the normal mechanical properties of the IVD in *ATC;Adgrg6^f/f^* mutant mice.

**Figure 3:**
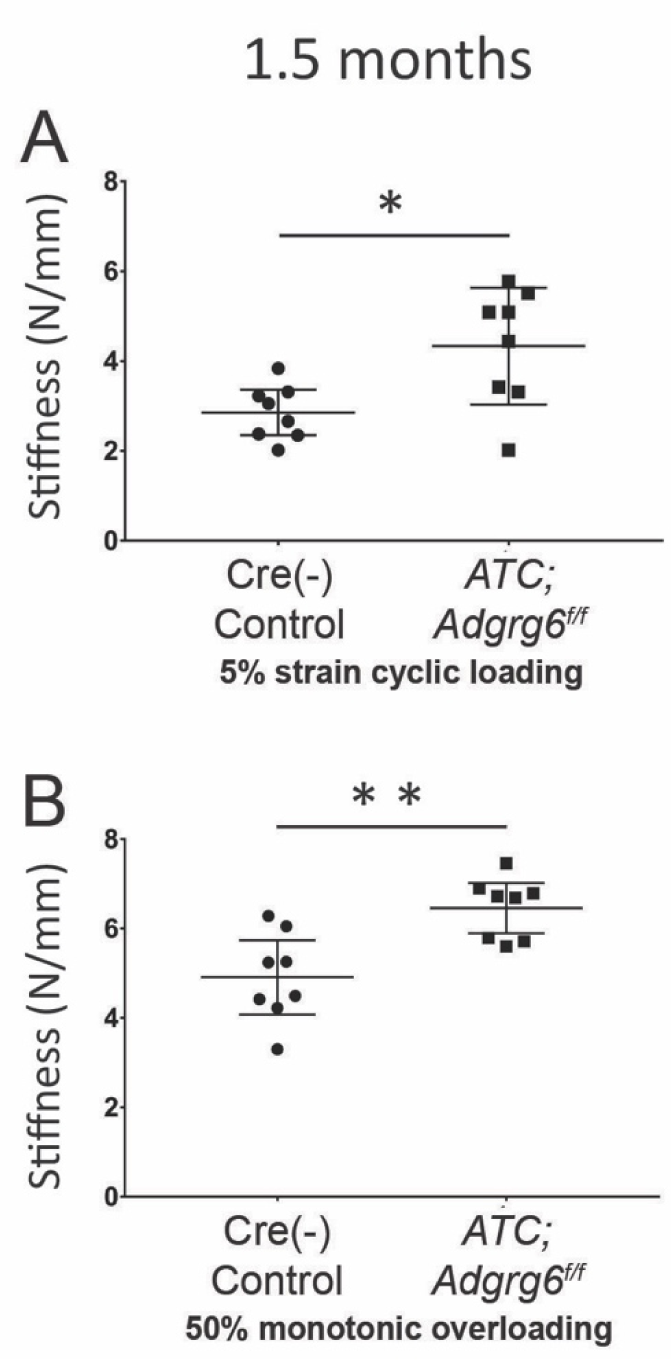
Young *ATC;Adgrg6^f/f^* mutant mice display increased mechanical stiffness of the IVD. (A, B) Mechanical testing using 5% strain cyclic loading (stiffness mean w/95% CI, p < 0.05) (A), and 50% monotonic overloading (stiffness mean w/95% CI, p < 0.01) (B), demonstrating increased stiffness in *ATC;Adgrg^f/f^* mutant lumbar IVDs. (*n*= 4 for each group, 4 IVDs were analyzed /mouse.)

### Loss of ADGRG6 in the intervertebral disc leads to increases in fibrotic extracellular matrix genes and alterations in ion transport components

To obtain additional, unbiased insights of the molecular and cellular changes in *Adgrg6* deficient mice prior to overt disc degeneration, we applied transcriptomic analysis on IVDs of P20 *Adgrg6*-defeicient mice. To avoid the contamination of untargeted IVD tissue in the *ATC;Adgrg6^f/f^* mutant mice (e.g. the outer most annulus fibrosus, Supplemental Figure 3C, D), we choose to use *Col2Cre;Adgrg6^f/f^* mutant mice which completely recombines throughout the entire IVD for these studies (Supplemental Figure 3B, E). Importantly, we observe analogous disc herniations in 8-month-old *Col2Cre;Adgrg6^f/f^* mutant mice (Supplemental Figure 7). We generated three independent libraries for each genotype (*Col2Cre;Adgrg6^f/f^* and Cre (-) control) from extracted IVD tissues (T8/9-L4/5), pooled from 2-3 individual mice at P20. We found 884 differential expressed genes with statistical significance (*p* value < 0.05) (Supplementary Table 1), and with a more stringent cut-off adjusted *p* value <0.05 and fold-change ≥2, we observed 42 differential expressed genes (Figure 4A). Enriched pathways and biological processes using gene ontology (GO) terms (Huang da et al., 2009) included extracellular matrix, positive regulation of fibroblast proliferation, extracellular matrix structural constituent conferring tensile strength, regulation of tyrosine phosphorylation of STAT protein, and ion transport (Figure 4B and Supplementary Table 2). Several of the significantly upregulated genes are associated with biomarkers or risk of lumbar disc degeneration and osteoarthritis in humans or animal models, including *Aspn*, *Dkk-3*, and *Mmp3* (Balakrishnan et al., 2014; Kizawa et al., 2005; Mahr et al., 2006; Snelling et al., 2016; Zhang et al., 2016) (Supplementary Table 1). We also observed alterations of chondrogenic and catabolic gene expression (Figure 4C, D). However, surprisingly some of the genes altered in *ATC;Adgrg6^f/f^* mutant mice at 1.5 months by qPCR analysis (Figure 2O), such as *Sox9, Col2a1* and *Mmp13,* were not similarly changed in slightly younger mice *Col2Cre;Adgrg6^f/f^* mutant mice (P20) by RNA-Seq analysis. However, in both conditional mutant mice we did observe a consistent upregulation of several fibrillar type collagens, including *Col1a1*, *Col3a1* among others at P20 in *Col2Cre;Adgrg6^f/f^* mutant mice (Figure 4E) and *Col1a1* and *Col3a1*in *ATC;Adgrg6^f/f^* mutant mice at 1.5 months (Figure 2O). Similar shifts from non-fibrous, predominantly type II collagens to fibrillar collagens, such as type I, are common signals reported in studies of degenerative IVD from both human and mouse models (Antoniou et al., 1996; Chen et al., 2016; Yee et al., 2016; Zhang et al., 2018). We suggest that the increased expression of more fibrotic extracellular matrix genes may underlie increased stiffness of the IVD we observed in *ATC;Adgrg6^f/f^* mutant mice (Figure 3) and contribute to susceptibility of endplate-oriented disc herniations during aging in these mutant mice (Figure 1).

**Figure 4:**
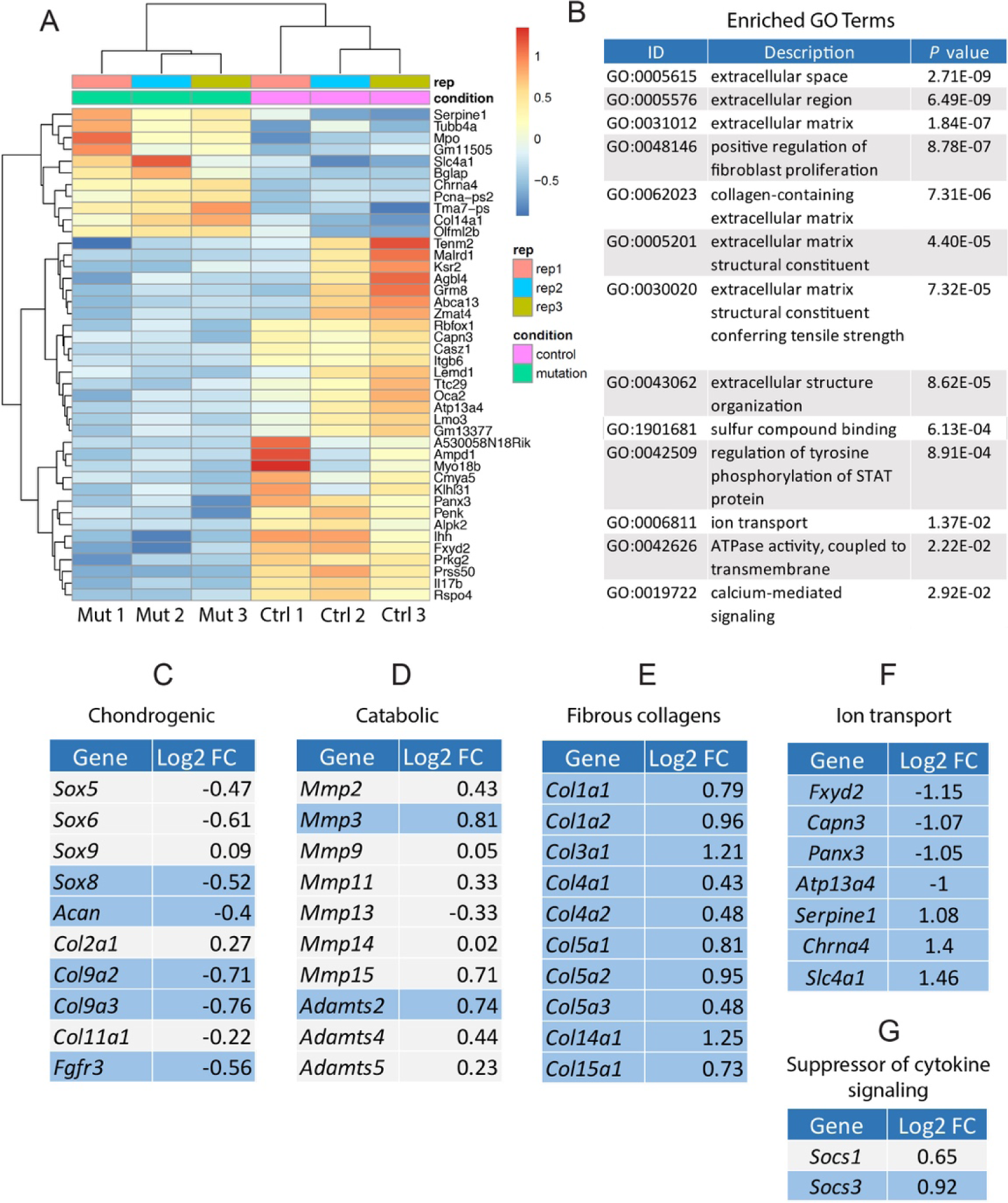
Young *Col2Cre;Adgrg6^f/f^* mutant mice display fibrotic-like changes of gene expression and dysregulation of genes associated with ion transport in the IVD. (A) Heatmap of differentially expressed gene based on RNA-sequencing analysis of IVDs derived from both *Col2Cre;Adgrg^f/f^* mutant (Mut 1-3) and Cre (-) littermates (Ctrl 1-3) at P20. (B) Gene ontology (GO) analysis revealed a suite of differentially expressed genes important for extracellular matrix organization and ion transport. (C-G) RNA-sequencing analysis revealed mild alterations in some chondrogenic (C) and catabolic (D) gene expression, but significantly induced fibrotic gene expression (E) and dysregulation of genes involved in ion transport (F) in the *Col2Cre;Adgrg^f/f^* mutant IVDs at P20. Some genes encode members of the suppressor of cytokine signaling were also upregulated in the mutant IVDs (G). Differential expressed genes with *p* value < 0.05 were highlighted in blue.

In addition, we observed gene expression changes in several genes associated with ion transport in the IVD of *Col2Cre;Adgrg6^f/f^* mutant mice at P20, including reduced expression of *Fxyd2*, *Capn3*, *Panx3*, and *Atp13a4*, as well as increased expression of *Serpine1*, *Chrna4*, and *Slc4a1* (Figure 4F). High osmotic pressure is a characteristic of the IVD (Fearing et al., 2018) and ion channel activity plays a critical role in the regulation of osmotic changes (Erickson et al., 2001; Matta et al., 2015). In agreement, recent analysis of the SM/J isotype mouse model of disc degeneration mice is associated with gene expression changes in ion transport systems (Zhang et al., 2018). Altogether, our transcriptomic analysis of the IVDs in *Col2Cre;Adgrg6^f/f^* mutant mice demonstrated a robust dysregulation of several important pathways and components of the IVD homeostasis, including induction of fibrotic gene expression, alteration of ion transport components, as well as changes in some chondrogenic and catabolic factors, prior to the onset of histopathology and disc degeneration. These data strongly suggest that ADGRG6 signaling is a critical regulator of postnatal homeostatic processes of the IVD.

### ADGRG6 regulates STAT3 signaling in multiple cartilaginous lineages

Our RNA-Seq analyses also implicate pro-inflammatory signaling involved in ADGRG6-deficient IVDs. For example, the *suppressor of cytokine signaling* (*Socs*) genes, *Socs1* and *Socs3* were significantly increased in *Col2Cre;Adgrg6^f/f^* mutant mice (Figure 4G) and implicate pathways associated with inflammation and activation/phosphorylation of STAT proteins (Figure 4B). SOCS3 directly regulates STAT1 (signal transducer and activator of transcription 1) and STAT3 activation (Carow and Rottenberg, 2014). For these reasons, we wanted to assay STAT1 and STAT3 activity in the IVD of *ATC;Adgrg6^f/f^* mutant mice.

By IHC analysis, we observed a substantial increase in phosphorylated STAT3 (pSTAT3) signal, the active form of STAT3 protein in the endplate of the *ATC;Adgrg6^f/f^* mutant IVD at 1.5 months (Figure 5B, B’), prior to overt histopathology of the disc. Quantification analysis revealed that 16.7% of cells in the endplate of the mutant IVD are pSTAT3 positive (*n*=3 mice; 2-3 IVDs/mouse; *n*=526 cells total), in comparison to 5.2% of the control IVD (*n*=3 mice; 2-3 IVDs/mouse; *n*=417 cells total) (Figure 5C). Similar analysis using an antibody against pSTAT1 failed to detect any signal in either genotype (Figure 5D-F). Together these data suggest that ADGRG6 regulates STAT3 activation in endplate of the IVD.

**Figure 5:**
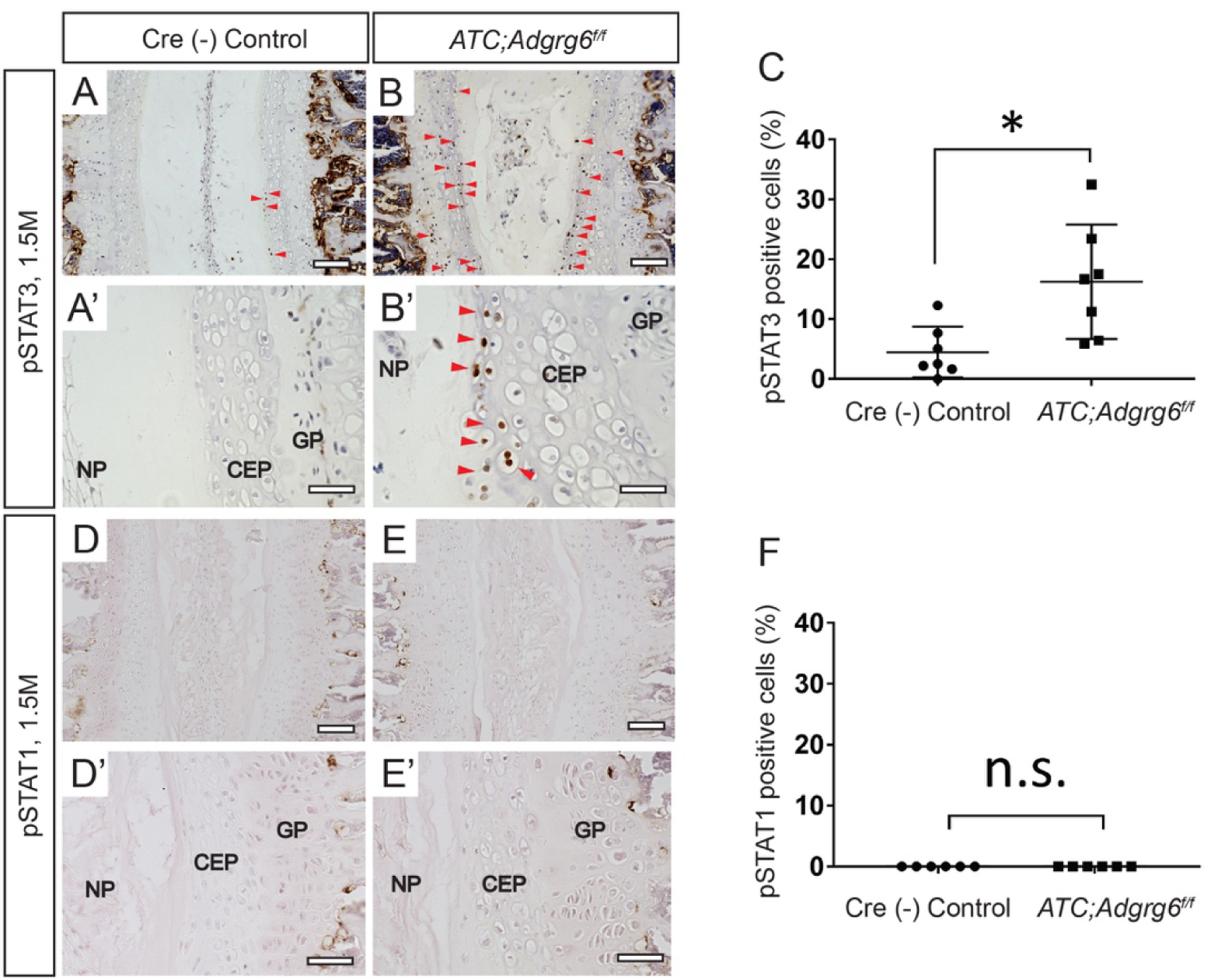
ADGRG6 regulates STAT3 signaling in IVDs. (A-B’) IHC analysis shows increased expression of pSTAT3 (red arrowheads, B, B’) in *ATC;Adgrg^f/f^* mutant mouse IVD at 1.5months (*n*= 3 for each group.) (C) Quantification of positive pSTAT3 cells in Cre (-) control or in *ATC;Adgrg^f/f^* mutant mouse IVDs. (*n*=3 mice for each group, at least two IVDs scored for each mouse. Dots plot with mean ± SD. *p≤0.05, Student’s *t* Test.) Scale bars: 200μm in (A, B), and 50μm in (A’, B’). *CEP-cartilaginous endplate, GP-growth plate, and NP-nucleus pulposus*.

After the induction of cytokines, including interleukin-6 (IL-6), IL-1, Tumor necrosis factor alpha (TNF), STAT3 is phosphorylated by receptor-associated Janus kinases and translocated to cell nucleus to regulate many cellular processes, such as cell growth and apoptosis (Garbers et al., 2015). Moreover, STAT3 activation is implicated for the progression of osteoarthritis (Hayashi et al., 2015; Latourte et al., 2017; Liu et al., 2015b). Our data indicated that ectopic, increased STAT3 activation occurs in young mice prior to histopathology of the mutant IVD, demonstrating a potential role of STAT3 signaling in initiation of endplate-oriented disc degeneration.

In order to facilitate more mechanistic studies of ADGRG6 function in the endplate and growth plate, we utilized the ATDC5 mouse cell line which can be induced to form cartilage-like tissue *in vitro* (Yao and Wang, 2013). We utilized CRISPR-Cas9 to disrupt the *Adgrg6* gene. After screening isolated clones, we identified a stable INDEL mutant with a homozygous 17-bp deletion in exon 3 of *Adgrg6* predicted to generate a frameshift mutation at amino acid Ser49 resulting in a truncated ADGRG6 protein (Adgrg6^p.Ser49+3fs^) (Figure 6A). The complete reduction of *Adgrg6* expression in our clonal *Adgrg6* mutant cell line (*Adgrg6* KO) suggested a null allele, likely due to non-sense mediated decay of the transcript (Figure 6B). During the course of chondrogenic maturation in unedited ATDC5 cells, *Adgrg6* expression increases with similar kinetics as other chondrogenic markers *Col2a1*, *Sox9* and *Acan* (Supplemental Figure 8). Consistent with our observations *in vivo* (Figure 2 C-N and O), we found decreased expression of several chondrogenic markers, including *Col2a1* and *Sox9,* and increased expression of hypertrophic marker (*Col10a1*) and catabolic enzyme (*Mmp13*) in these *Adgrg6* KO cells (Figure 6B). In addition, cleaved-Caspase 3, a key effector of apoptosis, was increased 2-fold in *Adgrg6* KO cells after 10-day maturation (Supplemental Figure 9), consistent with our observations of increased cell death in *ATC;Adgrg6^f/f^* mutant mice IVDs (Supplemental Figure 6). Consistent with the RNA-seq analysis of *Col2Cre;Adgrg6^f/f^* conditional mutant mice at P20, we also observed a 5-fold increase in the expression of *Socs3* in *Adgrg6* KO cells (Figure 6C), while *Socs1* was not detectable in either unedited wild-type or *Adgrg6* KO ATDC5 cells (data not shown). Taken together, the strong correlation between these *in vivo* and *in vitro* findings suggests a cell autonomous function for ADGRG6 in chondrogenic lineages for the regulation of typical gene expression profiles and of pro-inflammatory signals.

**Figure 6:**
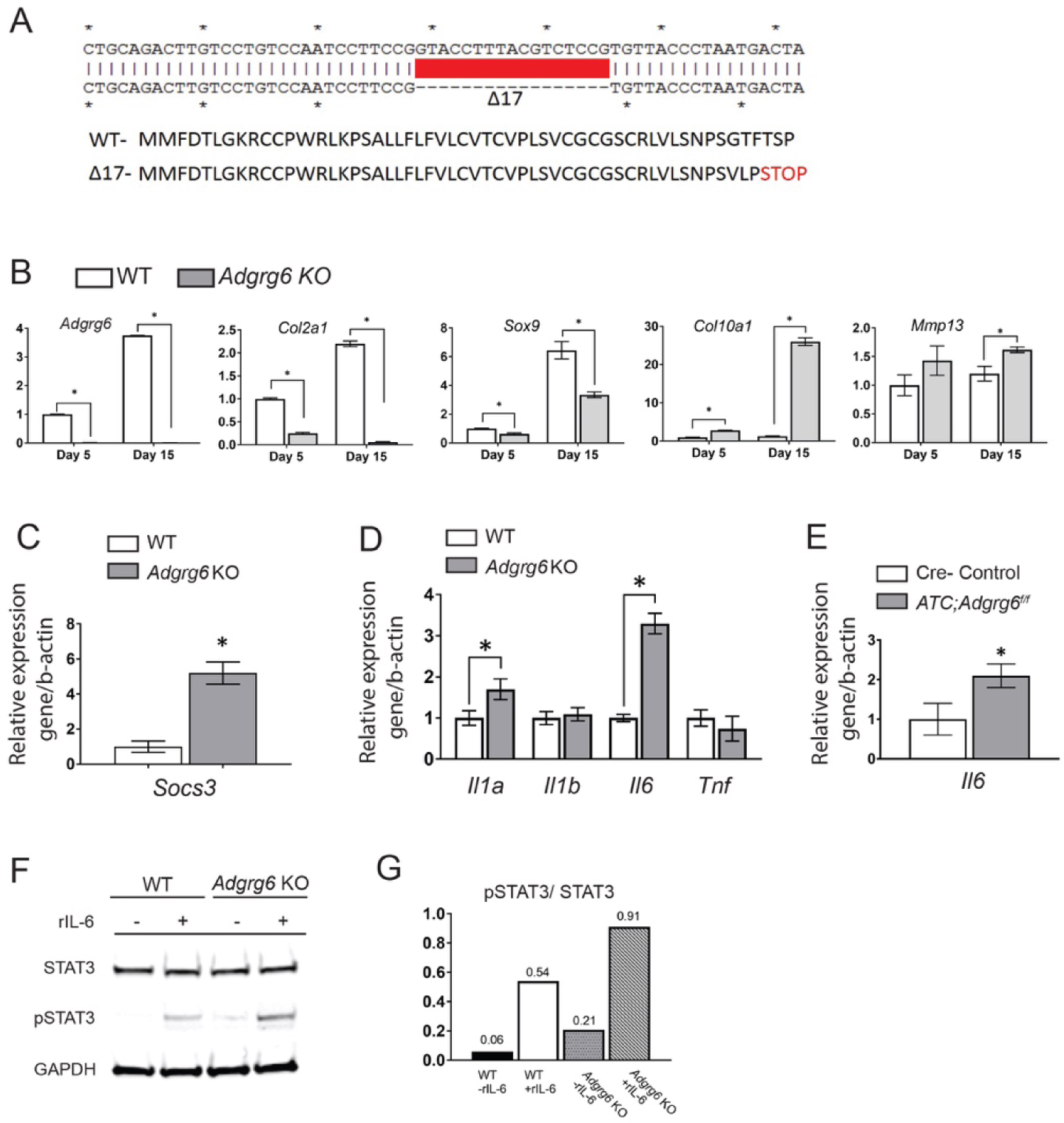
*Adgrg6* regulates gene expression profiles and STAT3 signaling in ATDC5 cell culture. (A) Schematic of a 17-bp deletion of *Adgrg6* from a stable single cell clone of ATDC5 cell line (*Adgrg6* KO). (B) Bright field image of ATDC5 cells maturated for 5 days demonstrates hypertrophic morphology in *Adrgr6* KO cells compared to wild type control cells. (C) qPCR analyses of gene expression in ATDC5 cells at 5 and 15 days of maturation demonstrates decreased expression of markers of healthy disc, *Col2a1*and *Sox9*, and increased expression of the hypertrophic marker, *Col10a1* and the extracellular matrix modifying enzyme, *Mmp13* in *Adgrg6* KO cells. (D) qPCR analyses revealed increased expression of *Socs3* in *Adrgr6* KO cells after 15 days of maturation. (E) qPCR analysis of *Il1a*, *Il1b*, *Il6* and *Tnf* in ATDC5 cells maturated for 15 days. (F) qPCR analysis of *Il6* expression in 1.5-month-old primary mouse IVDs. (C-F, *n*= 3 biological replicates and representative result is shown. Bars represent mean ± SD. *p≤0.05, Student’s *t* Test). (G, H) Representative Western blot and quantification of wild type and *Adgrg6* KO ATDC5 cell lysates showing stimulation of pSTAT3 staining after treatment with recombinant IL-6 (rIL-6) protein in both cell lines, while *Adgrg6* KO cells show a mild constitutive stimulation of pSTAT3 without addition of rIL-6 (*n*= 3 biological replicates and representative result is shown). Scale bars: 50μm in (B).

To better understand the intrinsic role of ADGRG6 in regulating STAT3 activation and the inflammatory signaling, we assayed a panel of known pro-inflammatory cytokines *Il1a*, *Il1b*, *Il6* and *Tnf* in *Adgrg6* KO cells (maturated for 15 days). We observed *Il6* expression was the most increased in *Adgrg6* KO cells (Figure 6D). Importantly, increased expression of *Il6* in the absence of ADGRG6 function was also observed in *ATC;Adgrg6^f/f^* mutant IVDs at 1.5 months of age (Figure 6E). IL-6 is the major upstream activators of STAT3 signaling (Garbers et al., 2015) that has been shown to associate with multiple degenerative joint diseases, including disc degeneration (Wuertz and Haglund, 2013), disc herniation (Eskola et al., 2012), as well as osteoarthritis (Livshits et al., 2009) in humans. IL-6 can not only stimulate the production of catabolic enzymes, but also can suppress the expression of anabolic genes, including *Sox9*, *Col2a1*, and *Acan* (Kapoor et al., 2011). To determine whether increased *Il6* expression was coupled with activation of STAT3 upon loss of ADGRG6 function, we assayed the ability of ATDC5 cells to respond to recombinant IL-6 protein (rIL-6) in culture. Western blot analysis demonstrated that rIL-6 effectively stimulates the expression of pSTAT3 in both unedited wild type and *Adgrg6* KO cells (Figure 6F, G). Interestingly, we detected constitutive pSTAT3 expression in *Adgrg6* KO cell lysates, which was further stimulated in expression by the addition of rIL-6 protein (Figure 6F, G). These *in vitro* results demonstrate that ADGRG6 regulates STAT3 expression and that IL-6 is a potential upstream activator of this signaling pathway in chondrogenic cell culture.

### STAT3 blockade protects against disc degeneration and herniation

To define the role of STAT3 signaling on the pathogenesis of endplate-oriented disc degeneration, we next tested if inhibition of STAT3 activation in the *Col2Cre;Adgrg6^f/f^* mutant mouse model which displays disc herniations by 8-months-of-age. To test this we utilized the constitutive *Col2Cre;Adgrg6^f/f^* conditional mutant in order to limit stress on the mice by avoiding exposure to Dox, as potential confounding variables, prior to long-term treatment with the STAT3 inhibitor, Stattic. Importantly, *Col2Cre;Adgrg6^f/f^* mutant mice exhibit similar phenotypes as observed in *ATC;Adgrg6^f/f^* mutant mice, including: erosion and clefts in the endplate and growth plate (yellow arrowheads; Figure 7B’ and supplemental Figure 8), ectopic expression of COLX in the endplate (red arrows; Figure 7E), increases in the appearance of endplate-oriented herniations with average of 9.6 herniations/mouse (*n*=3 mice; 39 IVDs) (Figure 7G). We treat both control and mutant groups with either a small molecule named Stattic, a nonpeptidic STAT3 inhibitor (25mg/kg, dissolved in DMSO), (Schust et al., 2006), or placebo (DMSO/PBS/Tween-20) via i.p. injection for 16 weeks beginning by the age of 1.5 months. Mutant mice receiving Stattic treatment displayed a reduction in endplate defects with an average of 3 herniations /mouse (*n*=3 mice; 39 IVDs) (Figure 7C, C’ and G). One way ANOVA followed by Tukey HSD test indicated that the mean score for the Stattic-treatment condition ((M)ean = 0.23, SD = 0.54) was significantly different (*p*<0.01) than the placebo-treated *Col2Cre;Adgrg6^f/f^* mutant mice (M = 0.74, SD = 0.56), but did not differ significantly from the placebo-treated Cre (-) control littermates (M = 0.096, SD = 0.09). Notably, placebo-treated Cre (-) control mice also display minor endplate defects with an average of 1.25 herniations/ mouse (*n*=4 mice; 52 IVDs), which is not observed in Cre (-) controls in Figure 1I. This may due to (i) DMSO content in the placebo treatment and (ii) different genetic background between *ATC;Adgrg6^f/f^* and *Col2Cre;Adgrg6^f/f^* mice. Of note, *Col2Cre;Adgrg6^f/f^* mutant mice receiving Stattic treatment displayed normal COLX expression in the endplate compared with the placebo-treated mutant mice (Figure 7E, F). In contrast, Stattic treatment had little effect on the general reduction of COLII and SOX9 expression in *Col2Cre;Adgrg6^f/f^* mutant mice (Supplemental Figure 10). These data suggest that additional effectors of ADGRG6 function, apart from STAT3 signaling, are required for maintenance of normal gene expression profiles in cartilaginous tissue of the IVD. In conclusion, these results suggest that blockade of STAT3 singling has a protective effect for endplate-oriented herniation, potentially through the repression of ectopic, increased COLX expression in the absence of ADGRG6 function.

**Figure 7:**
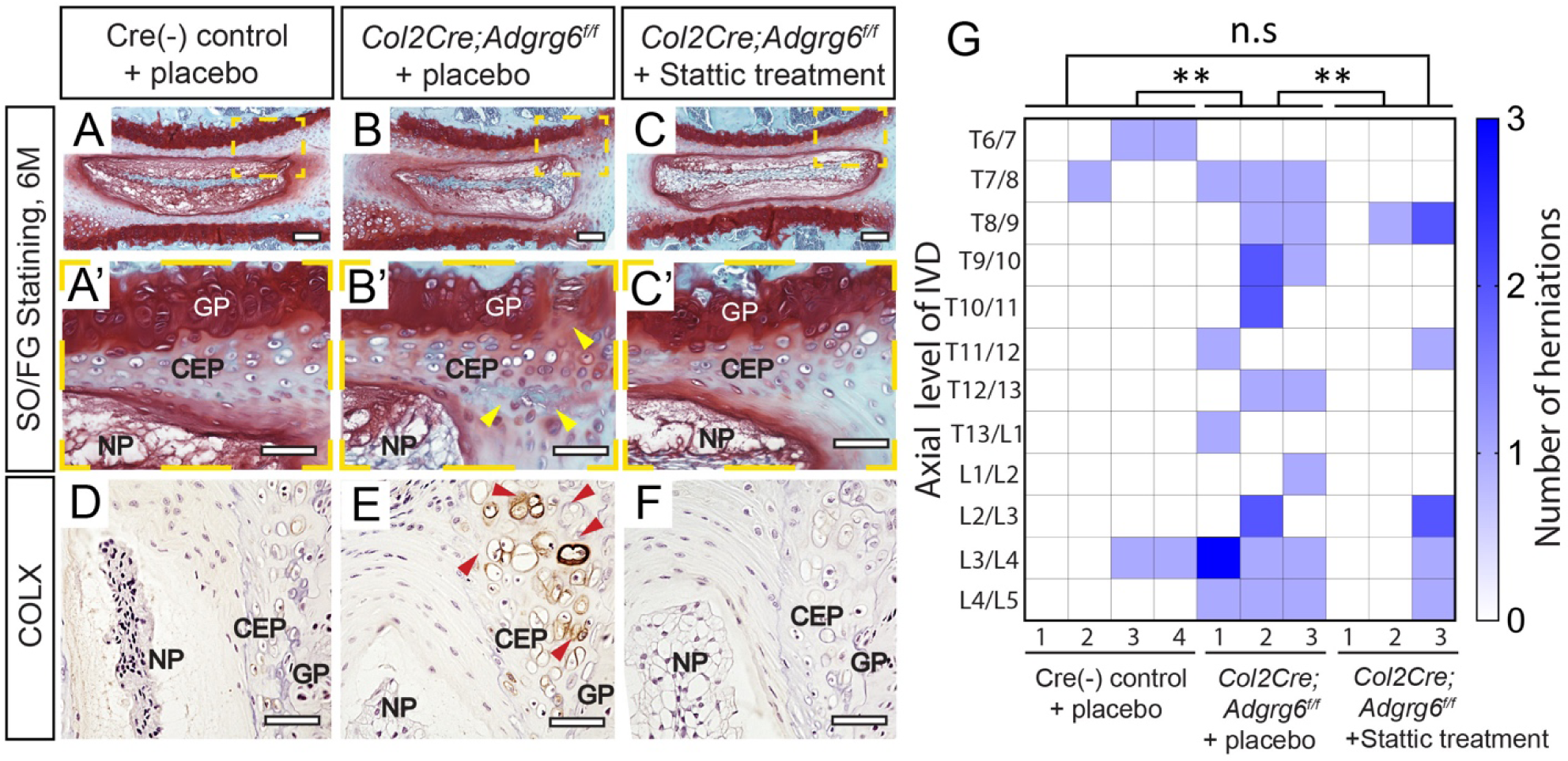
Inhibition of STAT3 by Stattic alleviates the formation of disc herniations attributed to loss of *Adgrg6* function. (A-C’) Representative Safranin-O/Fast green staining and (D-F) COLX IHC in medial-sectioned mouse IVD from placebo-treated Cre (-) control (A-A’, D), placebo-treated *Col2Cre;Adgrg6^f/f^* mutant (B-B’, E), and Stattic treated *Col2Cre;Adgrg6^f/f^* mutant mice (C-C’, F). *Col2Cre;Adgrg6^f/f^* mutant mice display defects of the IVD including lesions and clefts in the CEP and GP (yellow arrowheads, B’) and increased, ectopic expression of COLX within the CEP (red arrowheads, E), which is reduced by Stattic treatment (C-C’, F). (G) Heat map to represent contrast-enhanced microCT data from 6-month-old mice from three experimental groups: four placebo-treated Cre (-) controls; three placebo-treated *Col2Cre;Adgrg^f/f^* mutants; and three Stattic-treated *Col2Cre;Adgrg^f/f^* mutants. Plotted by the axial level of the IVD (left axis) and the number of herniations (right axis) observed in each mouse. (**p≤0.01, One way ANOVA followed by Tukey HSD test. n.s, not significant.) Scale bars: 200μm in (A-C), 50μm in (A’-C’) and (D-F). *CEP-cartilaginous endplate, GP-growth plate, and NP-nucleus pulposus*.

Taken together, our studies demonstrate that ADGRG6 has a cell autonomous role in the regulation of STAT3 signaling in chondrogenic lineages and with the IVD. Our demonstration that the small molecule STAT3 inhibitor Stattic can alleviate the onset and progression of degenerative changes and the formation of the endplate-oriented herniations in the IVDs of *Adgrg6*-deficient mice, implicating dysregulation of STAT3 signaling may underlie some forms of disc degeneration in particular for endplate-oriented disc degeneration. This implicates ADGRG6/STAT3 signaling pathway as a potential target for therapeutic approaches to treat disc degeneration, and possibly other joint degeneration disease, including osteoarthritis.

## Discussion

In this study, we demonstrate for the first time that ADGRG6 signaling acts as a positive regulator of postnatal homeostatic mechanisms of the IVD. This function is demonstrated by global changes in gene expression profiles in the IVD including alterations of several anabolic and catabolic factors and ion transport components, increased expression of fibrillar collagens and pro-inflammatory cytokine signaling, which occur several months prior to obvious histopathology of the disc in adult mice. We suggest that these alterations also lead to a general stiffening of the intervertebral disc, which synergistically contribute to the formation of endplate defects of the disc over the course of adult development.

In addition, we demonstrate that ADGRG6 acts as a cell autonomous regulator of the cytokine-induced STAT3 signaling in the cartilaginous endplate of the IVD, and that inhibition of this signaling provides protective effects against the onset and severity of endplate-oriented disc herniations due to the loss of ADGRG6 function. It will be interesting to test if blockade of STAT3 signaling has a general protective effect of endplate integrity that is not dependent on ADGRG6 function. However, to our knowledge there is not a well-established alternative experimental mouse model of endplate-oriented disc herniations with which to test this. It will be important to determine how STAT3 signaling may drive other models of disc degeneration characterized by lateral herniations and/or pathology of the nucleus pulposus.

The role of ADGRG6 appears to be dispensable for most developmental processes of cartilaginous tissues of the spine (Figure 2B). In contrast, we revealed a novel role for ADGRG6 function in the postnatal regulation of gene expression, stiffening of the IVD, and endplate-plate oriented disc defects in adult mice. Interestingly, we demonstrated that ADGRG6 is important for maintaining normal expression of SOX9, a master transcription factor for both chondrogenesis during embryonic development (Akiyama et al., 2002) and cartilage maintenance during postnatal development (Henry et al., 2012), in the IVD. This regulation of SOX/*Sox9* by of ADGRG6 function was recapitulated in heterologous chondrogenic ATDC5 cell culture, where we observed reduced expression of *Sox9* along with its direct target gene *Col2a1*. Conservation of this function is also observed in the cartilaginous semicircular canal of *adgrg6/gpr126* mutant zebrafish which similarly display altered expression of several extracellular matrix genes as well as decreased expression of *sox9b* (Geng et al., 2013). Furthermore, genetic ablation of *Sox9* in adult mice demonstrates reduction of *Adgrg6/Gpr126* expression, as well as many similarities as reported in this study including increased cell death and alterations of extracellular matrix gene expression (Henry et al., 2012). However, in contrast to our observations in *Adgrg6-*deficient mice, these *Sox9*-deficient mice also exhibit decreased disc height and loss of proteoglycan in the IVD (Henry et al., 2012). This may be explained by a more gradual loss of SOX9 expression in our conditional *Adgrg6* mouse model, which may stimulate unknown compensatory mechanisms that may overcome the onset of proteoglycan depletion observed in *Sox9-*deficent mice (El-Brolosy and Stainier, 2017). In addition, the ablation of *Adgrg6* in the IVD leads to unique endplate defects and alterations of a distinct suite of gene expression profiles, which was not reported after genetic ablation of *Sox9* in adult mice (Henry et al., 2012). Despite these observations, our study suggests a degree co-regulation exists for the expression of *Adgrgr6* and *Sox9*. The identification of the factors that govern this regulation and how this mechanistically contributes during homeostasis and disease warrants further investigation.

Our findings support a model where increased expression fibrillar collagens in IVD tissues in *Adgrg6* deficient mice precede the onset of increased disc stiffness and failure of the endplate. Using contrast-enhanced μCT imaging of the intact mouse spine, we for the first time describe the appearance and distribution of the endplate-oriented herniations in *Adgrg6* conditional mutant mice. Specifically, this mouse model of disc degeneration displays an upregulation of multiple fibrillar collagen genes (*e.g. Col1a1*, *Col3a1*, *Col4a1*, and *Col5a1*) in the IVDs at a juvenile stage of development (P20). Increased expression of fibrillar collagens and induced stiffness of the IVD is expected to result in disturbed stress distribution of the IVD, concentrate loading at the cartilaginous endplate, which increase the risk of endplate fractures and the formation of disc herniations (Adams et al., 1996; Vergroesen et al., 2015; Wilke et al., 2016). The adult IVD is thought to be an avascular tissue, as such its major source of nutrient flux occurs via diffusion through endplate (Urban et al., 2004). Increased expression of fibrillar collagens and catabolic enzymes in the disc including ectopic over-expression of COLX in the endplate are correlated with endplate sclerosis and reduced nutrient supply (Smith et al., 2011; Zhao et al., 2007). In this way a viscous cycle comprising increased expression of catabolic enzyme activity, pro-inflammatory factors, and apoptosis, can further exacerbate the integrity of the IVD leading to endplate-oriented herniation (Vergroesen et al., 2015). In conclusion, we suggest that increased fibrillary collagen expression and stiffness of the disc, increased ectopic expression of COLX in the endplate coupled with increases other catabolic factors such as MMPs and ADAMTSs, synergistically contribute to the endplate-specific defects in this mouse model.

Our previous work demonstrated a clear role for *Adgrg6* in the formation of scoliosis in mouse (onset at around P20-P40) (Karner et al., 2015). Here we show additional defects of endplate-oriented disc herniations in both *Col2Cre*- and *ATC;Adgrg6^f/f^* mutant mouse models (onset at around 6 to 8 months of age), which are reminiscent of an idiopathic human condition called acute Schmorl’s nodes (Adams and Dolan, 2012). While increased co-occurrence of Schmorl’s nodes in idiopathic scoliosis patients has been described (Buttermann and Mullin, 2008), additional studies in a larger cohort of AIS patients are needed to understand if an association between these IVD pathologies is relevant in the pathogenesis of disease in humans. Interestingly, while the incidence of scoliosis in *Col2Cre-* and *ATC;Adgrg6^f/f^* mutant mice was ∼80% and ∼12% respectively, herniations were observed in ∼100% of mutant mice in both models (Liu and Gray, unpublished data, Figure 1I and Figure 7G). Scoliosis in these mutant mice is specifically associated with curvature in the thoracic spine, while endplate-oriented herniations were observed along the entire spine axis. Taken together, our observations suggest that the mechanisms which promote the formation of endplate-oriented disc herniations in adult mice are partially independent from the pathogenesis of postnatal-onset idiopathic scoliosis in these mutant mice. It is tempting to speculate that alterations of the mechanical properties of the IVD during postnatal development underlie the susceptibility of scoliosis at this stage of development. However, additional and ongoing studies are required to better understand the cellular pathogenesis underlying scoliosis in *Adgrg6* mutant mice.

We demonstrate an upregulation of IL-6/STAT3 signaling, coupled with increased *Socs3* expression in *Adgrg6* conditional mutant mice, prior to overt histopathology. Furthermore, analysis of *Adgrg6*-KO ATDC5 cells in culture show activation of the IL-6/STAT3/*Soc3* pathway is an intrinsic property of cartilaginous cells, regulated by ADGRG6 function. Interestingly, IL-6/STAT3 signaling has been demonstrated to promote chondrogenic differentiation of human mesenchymal stem cells (Kondo et al., 2015), on the other hand IL-6/STAT3 signaling has been implicated in both degenerative disc disease and osteoarthritis. For example, induced expression of IL-6 and activation of STAT3 (pSTAT3) has been observed in human patients with disc degeneration and disc herniation (Osuka et al., 2014; Suzuki et al., 2017). Moreover, circulating IL-6 is positively associated with radiographic osteoarthritis and cartilage loss of the knee in humans (Stannus et al., 2010), and IL-6/STAT3 signaling is activated in trauma-induced osteoarthritis (Latourte et al., 2017; Liu et al., 2015b). IL-6 can also stimulate the expression of catabolic markers, such as COLX and MMP13 which are commonly associated with degenerative joint disease (He et al., 2014; Hunter et al., 2014). Importantly, systemic inhibition of STAT3 signaling with Stattic (Schust et al., 2006) can alleviate the onset and progression of joint remodeling in a post-traumatic osteoarthritis model in mouse (Latourte et al., 2017). However, how STAT3 activation is related to the initiation and progressions of these joint degenerative disorders remains to be determined. Nevertheless, we were able to demonstrate the alleviation of some aspects of endplate-oriented disc degeneration caused by *Adgrg6*-deficiency in the IVD with STAT3 inhibitor (Stattic) treatment. Altogether, this study provides clear evidence of the therapeutic value of STAT3 signaling for the onset and progression of disc degeneration.

Our work of comprehensive analysis of IVD development with both embryonic and postnatal deletion of *Adgrg6* also provide the first glimpse into the function and signal properties of an adhesion G-Protein coupled receptor (aGPCR) in this process. aGPCRs can function as classical GPCRs to invoke G-protein dependent intracellular *cis* signaling (Hamann et al., 2015). Indeed, ADRGR6 signals through Gs-protein/cAMP (Liebscher et al., 2014; Mogha et al., 2013). Future efforts will seek to analysis of how stimulation of Gs-signaling may alleviate IVD pathology and cAMP responsive binding elements are associated with alterations of genes affected in these *Adgrg6* deficient models. Given that GPCRs are among the most druggable classes of proteins, our work presented here identify a potential therapeutic target for degenerative joint diseases, and should encourage the targeted survey of the *ADGRG6* locus in human cohorts of disc degeneration and osteoarthritis.

## Methods

### Mouse Strains

All animal studies and procedures were approved by the Animal Studies Committee at the University of Texas at Austin. Mice were housed in standard cages and maintained on a 12-hour light/dark cycle, with rodent chow and water available ad libitum. All mouse strains were described previously, including *Adgrg6^f/f^* (Taconic #TF0269) (Mogha et al., 2013); *Rosa26;LacZ* (B6;129S-*Gt(ROSA)26Sor*/J) (Soriano, 1999); *ATC* (Dy et al., 2012), and *Col2Cre* (Long et al., 2001). Doxycycline (Dox) was administered to *ATC; Adgrg6^f/f^* mice and littermate controls with two strategies: (i) inducing from embryonic day (E)0.5-postnatal day (P)20 by ad libitum feeding of Dox-chow (Test Diet, 1814469) to plugged isolated females, and supplemented with intraperitoneal (IP) injections of the pregnant dames once/week (10mg Dox/kg body weight) throughout the pregnancy until the pups were weaned at P20; (ii) inducting from P1-P20 by ad libitum feeding of Dox-chow to the mothers after the pups were born, and supplemented with intraperitoneal (IP) injections of the mothers once/week (10mg Dox/kg body weight) until the pups were weaned at P20. *ATC; Rosa-LacZ^f/+^* mice were induced with the same strategies. STAT3 inhibitor Stattic (25mg/kg, dissolved in DMSO) or placebo (DMSO/PBS/Tween-20) were administered to *Col2Cre; Adgrg6^f/f^* mutant mice or Cre (-) littermate controls via i.p. injection once/week for 16 weeks beginning by the age of 1.5 months. Mice were harvested at P2, P20, 1.5 months, 6 months and 8 months of age.

### Analyses of mice

Histological analysis was performed on thoracic spines fixed in 10% neutral-buffered formalin for 3 days at room temperature followed by 1-week decalcification in Formic Acid Bone Decalcifier (Immunocal, StatLab). After decalcification, bones were embedded in paraffin and sectioned at 5μm thickness. Safranin O/Fast Green (SO/FG) and Alcian blue Hematoxylin/Orange G (ABH/OG) staining were performed following standard protocols (Center for Musculoskeletal Research, University of Rochester). Immunohistochemical analyses were performed on paraffin sections with traditional antigen retrieval and colorimetric development methodologies with the following primary antibodies: anti-Collagen II (Thermo Scientific, MS235B), anti-Collagen X (Quartett, 1-CO097-05), anti-SOX9 (Santa Cruz Biotechnology, sc-20095), anti-Lubricin (PRG4) (Abcam, ab28484), anti-MMP-13 (Thermo Scientific, MS-825-P), anti-IL-6 (Abcam, ab6672), and anti-phospho-STAT3 (Cell Signaling, #9145). The Terminal deoxynucleotidyl transferase dUTP Nick-End Labeling (TUNEL) cell death assay was performed on paraffin sections with the In Situ Cell Death Detection Kit, Fluorescein (Roche) according to the manufacturer’s instructions. The b-galactosidase staining was performed on frozen sections as previously described (Liu et al., 2015b). Spines were harvested and fixed in 4% paraformaldehyde for 2 hours at 4 °C and decalcified with 14% EDTA at 4 °C for 1 week. Tissues were washed in sucrose gradient, embedded with Tissue-Tek OCT medium, snap-frozen in liquid nitrogen, and sectioned at 10μm with a Thermo Scientific HM 550 cryostat. *In situ* hybridization using a Digoxygenin-labeled antisense riboprobe for *Adgrg6* was performed on 5μm paraffin sections as described previously with modifications (Karner et al., 2015), and detected with either a chromogenic substrate (BM Purple, Roche) or a tyramine-amplified fluorescent antibody (Perkin Elmer).

### Cell culture

ATDC5 cells (Sigma, 99072806) were maintained in DMEM/F-12 (1:1) medium (Gibco, 11330032) supplemented with 5% FBS and 1% penicillin/streptomycin. ATDC5 cells were differentiated in DMEM/F-12 (1:1) medium supplemented with 5% FBS, 1% penicillin/streptomycin, 1% ITS premix (Corning, 354352), 50µg/ml ascorbic acid, 10nM dexamethasone, and 10ng/ml TGF-β3 (Sigma, SRP6552) for 5, 10, and 15 days. Both wild type and *Adgrg6* KO ATDC5 cells were treated with 100ng/ml recombinant human IL-6 protein (rIL-6) (R&D System, 206-IL) for 2 hours before protein extraction.

### Generation of the *Adgrg6* KO cell line

CRISPR reagents were generated to target the 3rd exon of mouse *Adgrg6* (ENSMUST00000041168.5) using the following oligos: *Adgrg6*-g33-fwd ACACCGAGGGTAACACGGAGACGTAAG and *Adgrg6*-g33-rev AAAACTTACGTCTCCGTGTTACCCTCG and cloned into a lentiviral packing vector (lentiCRISPR v2 was a gift from Feng Zhang (Addgene plasmid # 52961)) along a pCas9_GFP (a gift from Kiran Musunuru (Addgene plasmid # 44719)). Lentiviral particle packaging was in A293T cells using standard 3rd generation approach (https://tronolab.epfl.ch/page-148635-en.html). Human embryonic kidney (HEK) 293T cells (Sigma) were maintained in DMEM supplemented with 10% fetal bovine serum, 2mM GlutaMAX (Life Technologies), 100U/ml penicillin, and 100ug/mL streptomycin at 37°C with 5% CO_2_ incubation. 293T cells were seeded into 6cm plates (Corning) one day prior to transfection at a density of 2×10^6^ cells per well. 293T cells were transfected using FUGENE 6 (Promega) following the manufacturer’s recommended protocol. For each plate, a total of 0.5ug of each plasmid was used. At 2 and 4 days post transfection, the cell media was collected and filtered with 0.45 μM filter (Corning) and stored at −80°C.

ATDC5 were plated in 6-well plates to 80% confluency and lentiviral transduction was using diluted viral media with 0.1% polybrene (EMD Millipore) for 24 hours followed by selection with 4μg/ml Blastocidin and Puromycin for 5 days post transfection. Serial dilution under selection was used to identify individual clones, expanded colonies were screened for INDEL mutations using *Adgrg6*-ex3-fwd – TTGACAGTTACTGCTTGATGCCCCC and *Adgrg6*-ex3-rev- CCCTTGGCAGTCGCTCCACAGAATT primers and amplicons were screen by Sanger sequencing to identify homozygous clones.

### RNA isolation and qPCR

Intervertebral discs from the thoracic and lumbar spine (T8-L5) of the 1.5-month old *ATC; Adgrg6^f/f^* and control mice were isolated in cold PBS, snap frozen and pulverized in liquid nitrogen. Total RNA from intervertebral discs was isolated using the TRIzol Reagent (Invitrogen, 15596026), and cleaned up with the Direct-zol RNA miniprep kit (Zymo Research, Z2070). Total RNA of the cultured ATDC5 cells was isolated using the RNAeasy mini kit (Qiagen, 74104). Reverse transcription was performed using 1μg total RNA with the iScript cDNA synthesis kit (BioRad). Reactions were set up in technical and biological triplicates in a 96 well format on an BioRAD CFX96 real-time PCR detection system, using SYBR green chemistry (SsoAdvanced, BioRad). The PCR conditions were 95°C for 3 min followed by 40 cycles of 95°C for 10s and 58°C for 30s. Gene expression was normalized to *b-actin* mRNA and relative expression was calculated using the 2^−(ΔΔCt)^ method. All qPCR primers sequences are listed in Supplementary Table 3. PCR efficiency was optimized and melting curve analyses of products were performed to ensure reaction specificity.

### RNA isolation, library construction and sequencing

Intervertebral discs from the thoracic and lumbar spine (T8-L5) of the P20 *Col2Cre; Adgrg6^f/f^* and control mice were isolated in cold PBS, snap frozen and pulverized in liquid nitrogen. Total RNA was extracted using Trizol reagent (Invitrogen, CA, USA) following the manufacturer’s procedure. The total RNA quantity and purity were analyzed on Bioanalyzer 2100 and RNA 6000 Nano LabChip Kit (Agilent, CA, USA) with RIN number >7.0. Total RNA was subjected to isolate Poly (A) mRNA with poly-T oligo attached magnetic beads (Invitrogen). RNA fragments were reverse-transcribed to create the final cDNA libraries following the NEBNext® Ultra™ RNA Library Prep Kit (Illumina, San Diego, USA), paired-end sequencing was performed.

### Bioinformatics analysis

#### Transcripts Assembly

Firstly, Cutadapt (Martin, 2011) and perl scripts in house were used to remove the reads that contained adaptor contamination, low quality bases and undetermined bases. Then sequence quality was verified using FastQC (http://www.bioinformatics.babraham.ac.uk/projects/fastqc/). We used HISAT2 (Kim et al., 2015) to map reads to the genome of Mus Musculus (GRCm38.88). The mapped reads of each sample were assembled using StringTie (Frazee et al., 2015). Then, all transcriptomes from biological samples were merged to reconstruct a comprehensive transcriptome using perl scripts and gffcompare. After the final transcriptome was generated, StringTie (Pertea et al., 2015) and Ballgown (Frazee et al., 2015) was used to estimate the expression levels of all transcripts.

#### Different expression analysis of mRNAs

StringTie (Pertea et al., 2015) was used to perform expression level for mRNAs by calculating FPKM. The differentially expressed mRNAs were selected with log2 (fold change) >1 or log2 (fold change) <-1 and with statistical significance (p value < 0.05) by R package Ballgown (Frazee et al., 2015).

### Western blotting

For western blotting analysis, total proteins were extracted from cells with protein extraction buffer [50mM HEPES, 1.5mM EDTA (pH 8.0), 150mM NaCl, 10% glycerol, 1% Triton X-100] supplemented with protease and phosphatase inhibitors (Roche). 10mg of protein from each sample was resolved by 4-15% SDS-polyacrylamide gel electrophoresis and transferred to the nitrocellulose membranes. Western blots were then blocked with LI-COR blocking buffer and incubated overnight with primary antibodies anti-STAT3 (Cell Signaling, #4904), anti-pSTAT3 (Cell Signaling, #9145), and anti-GAPDH (Cell Signaling, #2118) at 4 °C with gentle rocking. The next day western blots were detected with the LI-COR Odyssey infrared imaging system.

### Contrast-enhanced μCT imaging and segmentation

Samples undergoing contrast-enhanced micro-computed tomography (μCT) were incubated in a 35%w/v solution Ioversol in PBS (OptiRay 350, Guerbet, St. Louis) supplemented with 1% penicillin–streptomycin at 37C one day prior to scanning. Immediately prior to scanning, the sample was removed from the solution and wrapped in PBS-soaked gauze. These samples were mounted in 2% agarose gels and then scanned using the microCT40 system (Scanco Medical, CH) operating at 6 μm voxel size (45kVp, 177uA, 300 ms integration). Following our previous method for segmentation of murine IVDs (Lin and Tang, 2017; Lin et al., 2016), the μCT CT data is exported as a DICOM file for further processing. Following an initial median filter (sigma = 0.8, support = 3), bone is then thresholded out, and the soft tissue not part of the IVD was removed by drawing contours around the outer edge of every five transverse slices of the AF and morphed using a linear interpolation. The remaining voxels are designated as the whole disc mask. From the masks of the whole disc, volumes and average attenuations (intensity) are calculated. The volume was determined from the total number of voxels contained within the mask and the attenuation is taken as the average 16-bit grayscale value of the voxels. Visualizations of the μCT were obtained using OsiriX (Pixmeo, Geneva). The volume of the contoured disc was then measured. Defects of the IVD resembling Schmorl’s Nodes were defined by three or more consecutive slices that had a rupture in the same area of the endplate. This method was chosen so as to eliminate any potential spatial artifacts that may be misidentified as ruptures.

### Mechanical testing

The mechanical properties of the isolated intervertebral discs were determined using dynamic compression on a microindentation system (BioDent 1000; Active Life Scientific, Santa Barbara, CA) with a 2.39 mm non-porous, flat probe (Liu et al., 2015a). The probe’s load cell resolution is 0.001 N, and the system’s Piezo actuator resolution is 0.01 μm. Each sample was moved into position under the probe tip by gripping the aluminum platen. The indenter tip was aligned over each sample so that the probe covered the entire diameter of the disc. Each disc was first loaded sinusoidally at 10% strain peak strain at 1 Hz for 20 cycles with a 0.1 N preload. After the cyclic tests, the discs were monotonically overloaded to 50% strain at a loading rate of 10% strain per second. The loading slope value was obtained from the linear region of the force displacement curve from all loading curves. These samples were maintained in physiological PBS solution (pH 7.2) during and between trials to simulate the osmotic pressures found in the body and maintain hydration of the IVD.

### Quantification of collagen and proteoglycans

The wet weight of each isolated disc was taken after mechanical testing utilizing an analytical balance (A-200DS; Denver Instrument Company, Bohemia, NY). Samples were first digested in papain at 65°C for 18 h. The samples were then centrifuged and the supernatant collected and then plated in triplicate. Proteoglycan content was quantified using the colorimetric dimethyl-methylene blue (DMMB) assay (Sabiston et al., 1985) by measuring the absorbance 525nm with chondroitin sulfate from bovine cartilage as standards (Sigma-Aldrich, St. Louis, MO), and then normalized to wet weight of the IVD. The remaining papain-digested lysates were then used for hyproxyproline quantification. The amount of collagen was approximated by assuming that hydroxyproline accounts 1/7 of the mass of collagen. The samples were hydrolyzed in 12 N hydrochloric acid at 120 °C for 3 h. The hydrolyzed samples were then plated in triplicates. A chloramine T colorimetric assay (Woessner, 1961) and standardized using hydroxyproline by quantifying the absorbance at 560 nm using a plate reader (SpectraMax M2, Molecular Devices, Sunnyvale, CA). Quantification of proteoglycan content in ATDC5 cell culture were conducted as previously described with modifications (Kitaoka et al., 2001). Briefly, ATDC5 cells were fixed with 10% neutral buffered formalin for 10 minutes at room temperature and incubated with 3% acetic acid for 10 minute, followed by staining with 1% Alcian blue (in 3% acetic acid, pH 2.5) for 30 minutes at room temperature. Cells were washed with PBS and air-dried overnight. After taking pictures, Alcian blue was extracted with 500ul of DMSO and quantifying the absorbance at 650nm using a plate reader. At least three biological replicates were analyzed for each experimental group.

### Statistics

Statistical analyses to compared the mutant and control groups were performed using 2-tailed Student’s *t*-test and one-way ANOVA followed by Turkey HSD test as appropriate (GraphPad Prism 7). A *p* value of less than 0.05 is considered statistically significant.

## Author contributions

ZL and RSG conceived the experiments. ZL, GE, and NM performed the experiments. ZL, NM, and JZ performed RNA-sequencing and analysis. RSG, SYT, NM, NA and MJH provided reagents, expertise, and feedback. ZL and RSG wrote the manuscript.

## Acknowledgements

We would like to thank Dr. Fanxin Long and Dr. Veronique Levebre for sharing the *Col2Cre* and *ATC* mouse strains respectively. We would like to thank Drs. John Wallingford and Steve Vokes, and members of the Gray lab for helpful comments on this manuscript prior to submission. Research reported in this publication was supported by the National Institute of Arthritis and Musculoskeletal and Skin Diseases of the National Institutes of Health under Award Number R01AR072009-01 (R.S.G.), F32AR073648 (Z.L.), R21AR069804 (S.T.), R01AR063071 (M.J.H) and the National Institute of Child & Human Development 1P01HD084387 (N.A).

## Supplemental Figures

**Supplemental Figure 1:**
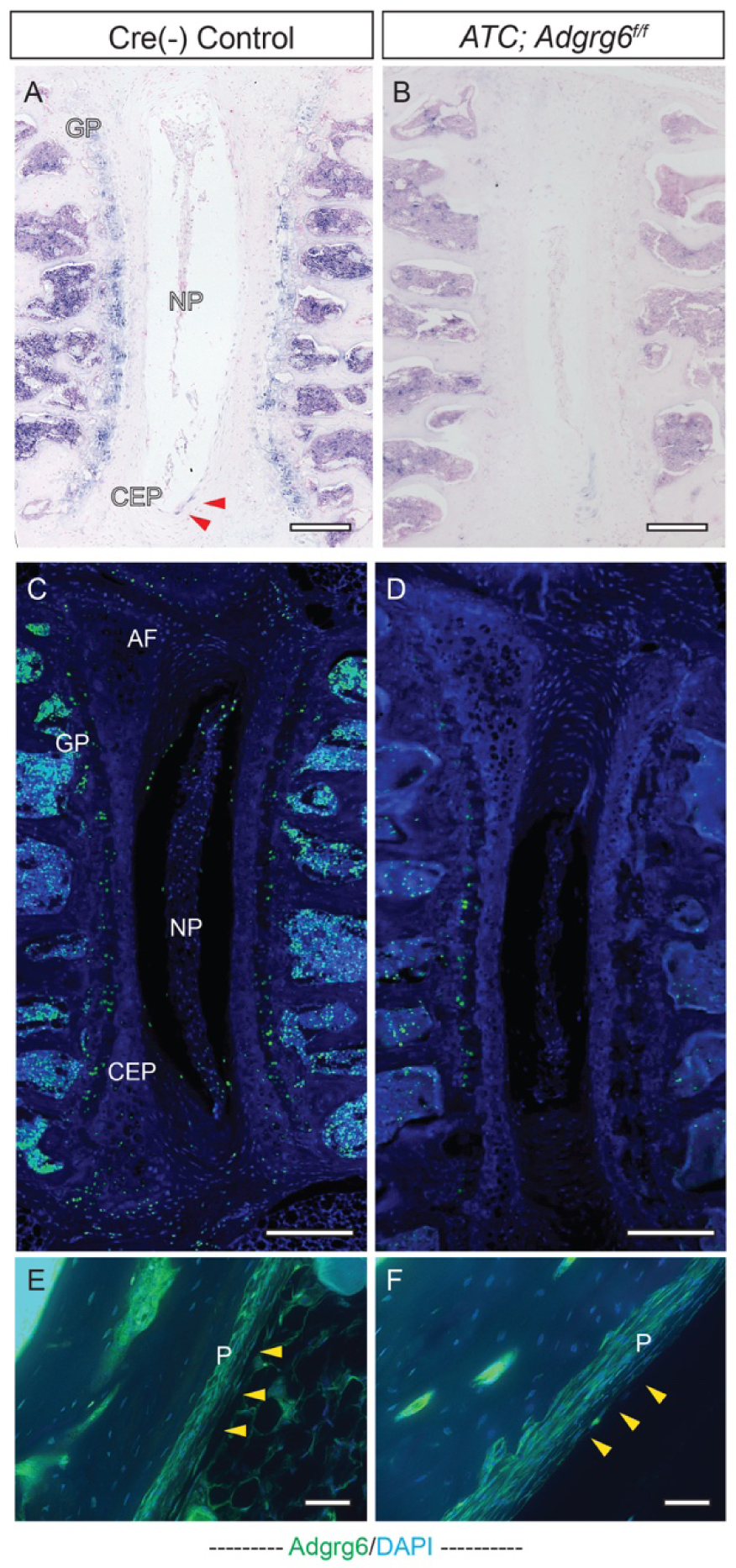
*In situ* expression of *Adgrg6* in the spine. (A, B) *In situ* hybridizations of *Adgrg6* in spine tissue (8 months) using Alkaline phosphatase/BM purple chromogenic developing shows strong *Adgrg6* expression (blue stain) in the growth plate (GP) and minor expression in the annulus fibrosis (AF) (red arrowheads) that is mostly abolished in *ATC;Adgrg6^f/f^* mutant tissues (B); or using (C, D) tyramine-amplification fluorescence which shows expanded expression throughout the IVD including GP, CEP, AF, and NP, which is mostly diminished in *ATC;Adgrg6^f/f^* mutant tissues. Robust expression was detected in periosteum of the long bone tissues in both control and the *ATC;Adgrg6^f/f^* mutant moues (E, F, yellow arrows). Scale bars: 200μm in (A-D); 50μm in (E, F). *CEP-cartilaginous endplate, GP-*

**Supplemental Figure 2:**
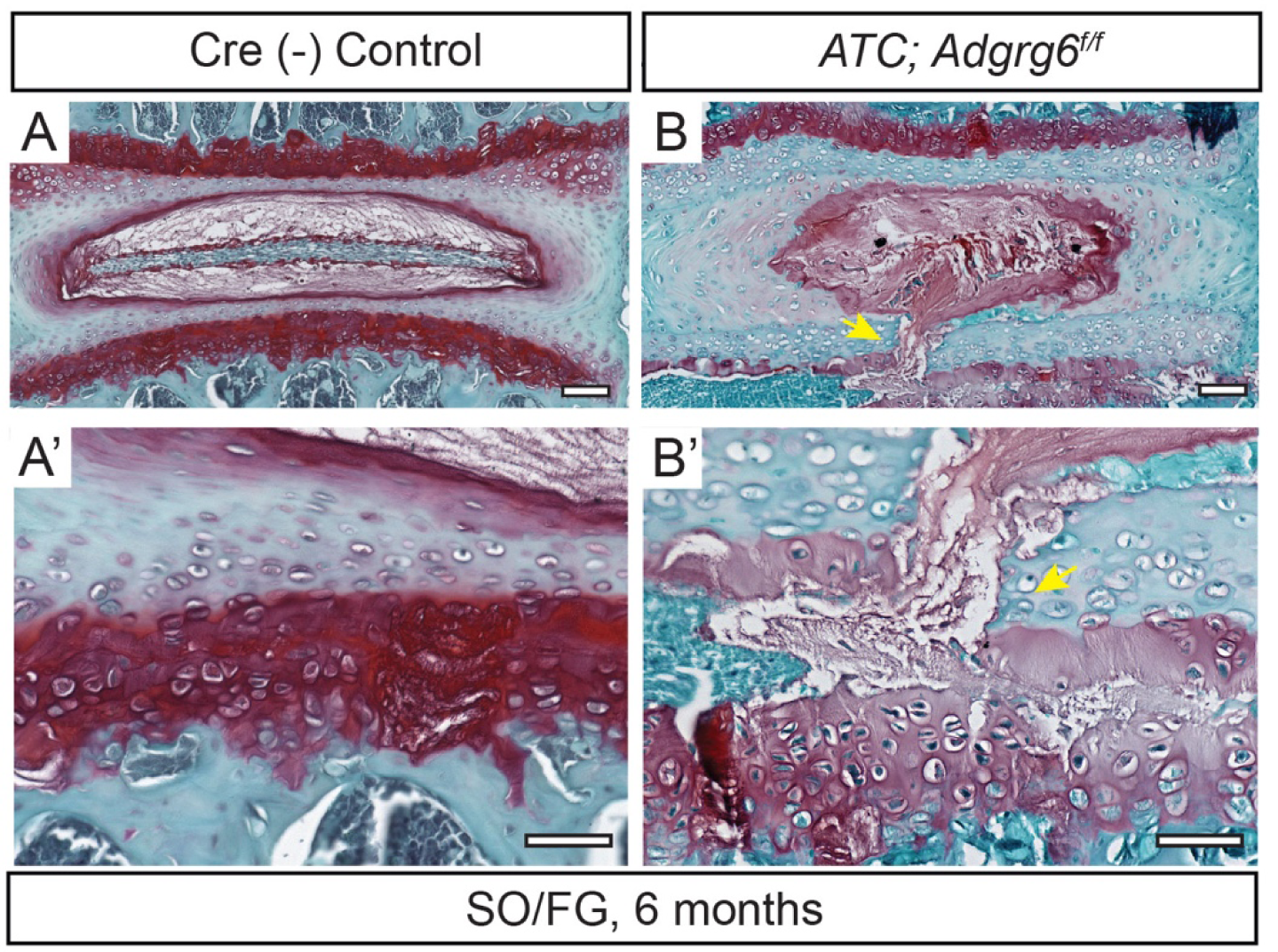
Embryonic loss of Adgrg6 lead to endplate-oriented herniations of the IVD in adult *ATC;Adgrg6^f/f^* mutant mice. (A-B’) Representative medial-sectioned 6-month-old mouse IVDs (induced form E0.5-P20) stained with Safranin-O/Fast green (SO/FG) (*n*=3 for controls and *n*=6 for mutants). Endplate-oriented disc herniation is indicated with yellow arrows. Scale bars: 100μm in (A, B); 50μm in (A’, B’).

**Supplemental Figure 3:**
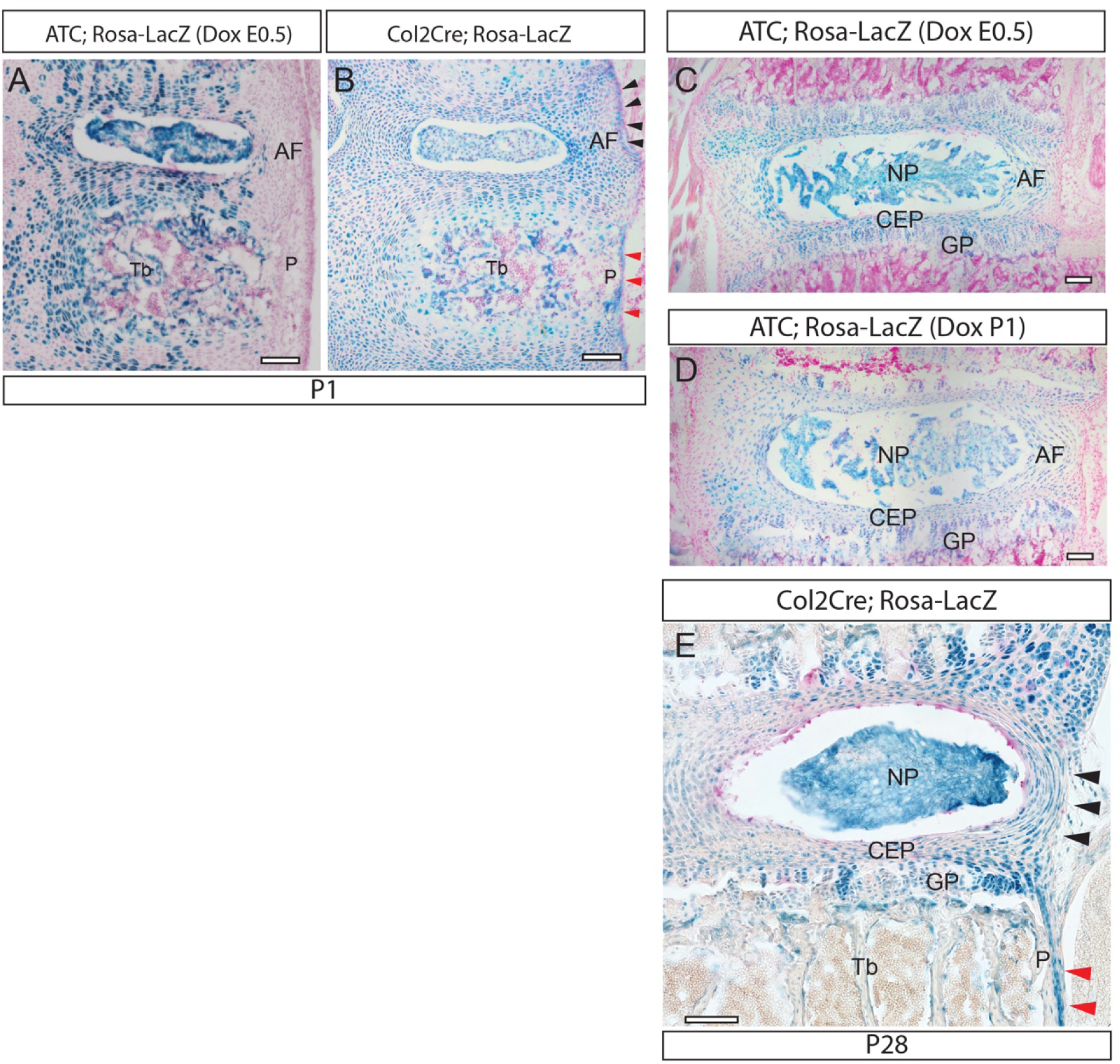
β-galactosidase staining of LacZ-reporter mice induced with different strategies. More robust recombination (blue signal) in CEP, GP, and AF of the IVD was observed in the *Col2Cre; Rosa-LacZ* mouse (B, E) compared with the *ATC; Rosa-LacZ* mouse when induced from E0.5-P20 (A, C) and P1-P20 (D). Recombination in periosteum (B, E, red arrows) and the outmost AF layers of the IVD (B, E, black arrows) was observed only in the *Col2Cre; Rosa-LacZ* mouse but not the *ATC; Rosa-LacZ* mouse. Scale bars: 100μm in (A-E). *CEP-cartilaginous endplate, GP-growth plate, AF-annulus fibrosis, NP-nucleus pulposus, Tb-trabecular bone, and P-periosteum*.

**Supplemental Figure 4:**
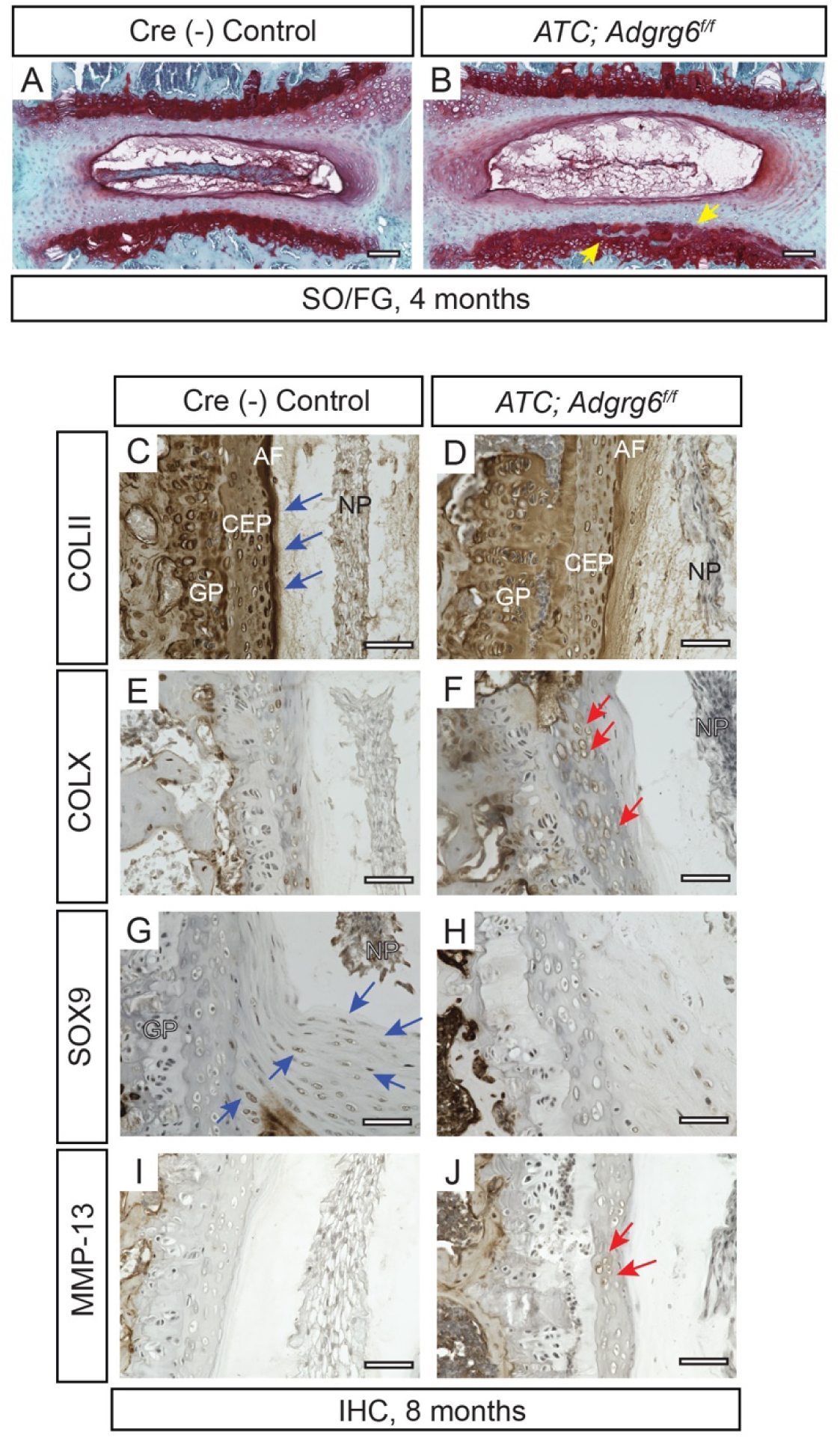
Postnatal loss of *Adgrg6* in *ATC;Adgrg6^f/f^* mutant mice leads to degenerative changes in the IVDs. (A, B) Representative medial-sectioned 4-month-old mouse IVDs stained with Safranin-O/Fast green (SO/FG) (induced from P1-P20, *n*=3 for controls and *n*=5 for mutants). Minor growth plate erosion is observed in two out of five mutant mice (B, yellow arrowheads). (C-J) IHC analysis of 8-month-old Cre (-) Control and *ATC;Adgrg^f/f^* mutant mouse IVDs (induced from P1-P20). Several protein markers of IVD health and disease are affected in *ATC;Adgrg^f/f^* mutant IVD including decreased expression of healthy disc markers COLII (C, blue arrows) and SOX9 (G, blue arrows), and increased expression of the hypertrophic marker COLX (F, red arrows) and extracellular matrix modifying enzyme MMP-13 (J, red arrows). (*n*=3 for each group.) Scale bars: 100μm in (A, B); 50μm in (C-J). *CEP: cartilaginous endplate, GP: growth plate, AF: annulus fibrosis, and NP: nucleus pulposus*.

**Supplemental Figure 5:**
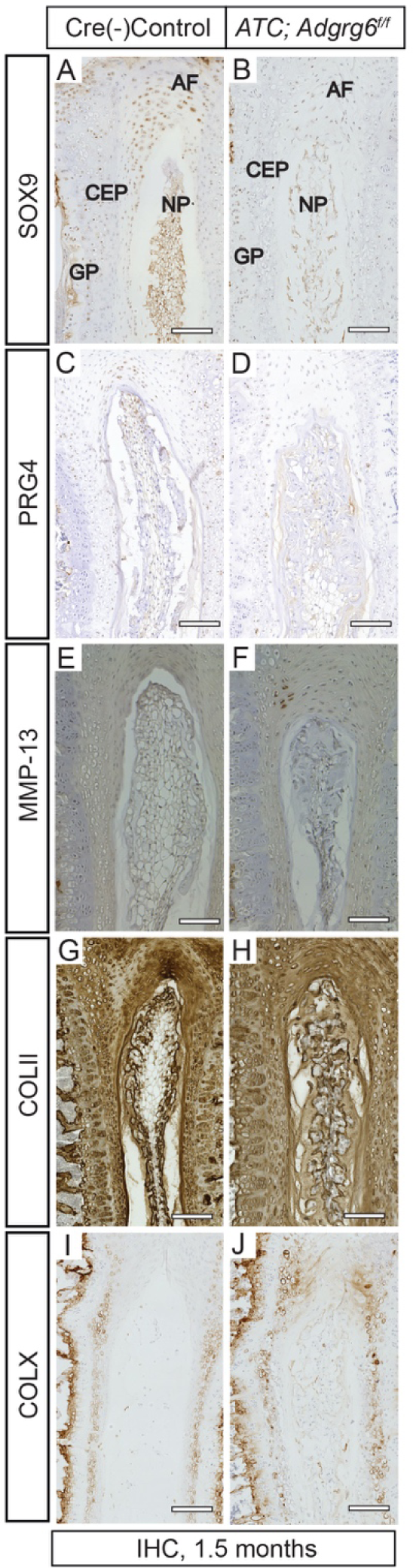
Young *ATC;Adgrg6^f/f^* mutant mice display degenerative alterations of protein expression in the IVD. Large scale images of IHC analysis shown in Figure 2. IHC analysis of common markers of degenerative disc. *ATC;Adgrg^f/f^* conditional mutant IVDs display reduced expression of markers of healthy disc: SOX9 (B), PRG4 (D), and COLII (H); and increased expression of extracellular matrix modifying enzymes MMP-13 (F), hypertrophic marker COLX (J). Scale bars: 100μm in (A-J). *AF-annulus fibrosis, CEP-cartilaginous endplate, GP-growth plate, and NP-nucleus pulposus*.

**Supplemental Figure 6:**
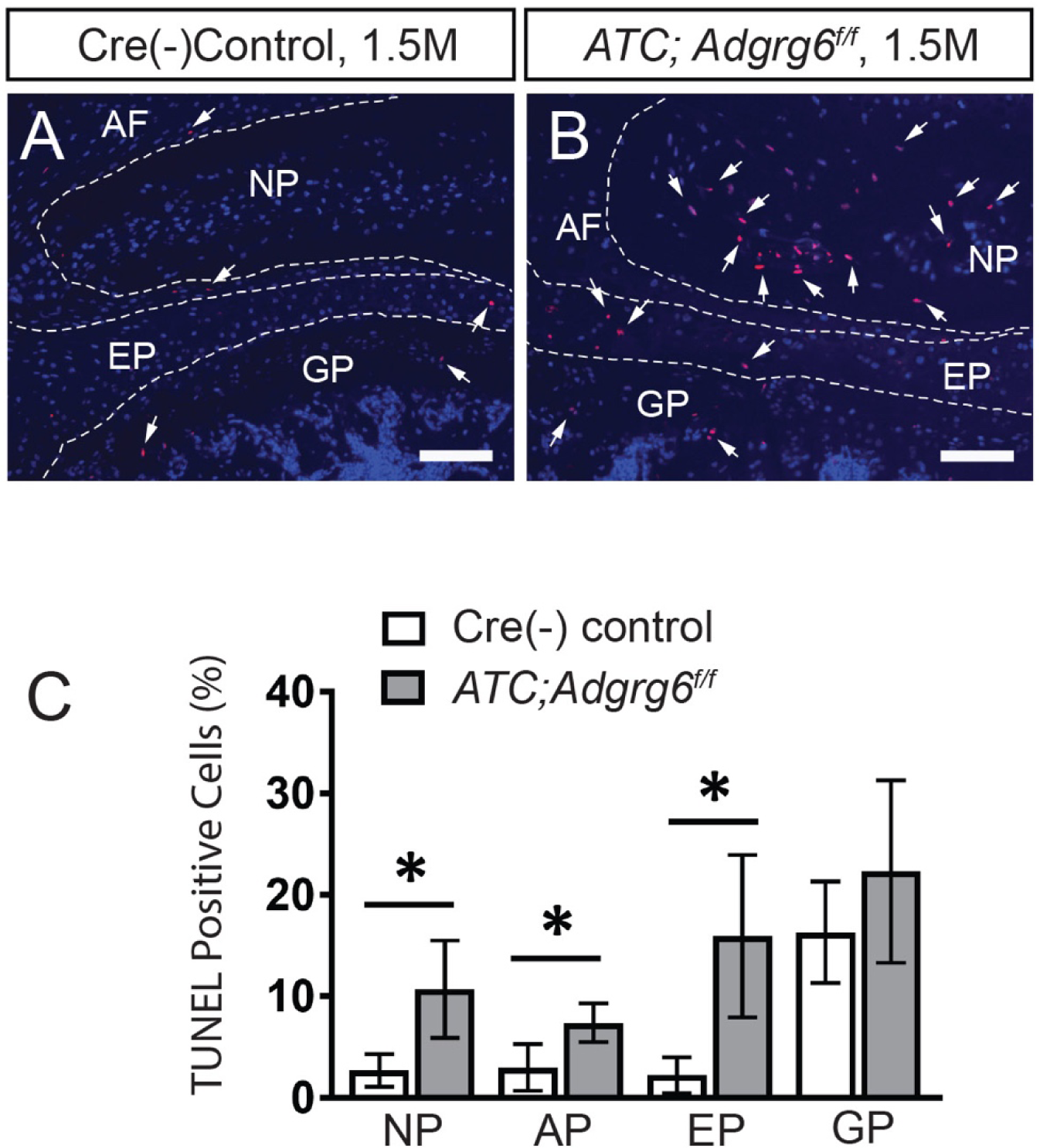
Young *ATC;Adgrg6^f/f^* conditional mutant mice display increased apoptosis in the IVD. (A, B) TUNEL (red fluorescence) staining of 1.5-month-old *ATC;Adgrg^f/f^* mutants (B, white arrows) display increased TUNEL positive cells compared to Cre (-) control (A) mice. (C) Graph of the ratio of TUNEL positive cells to total cells (DAPI) (*n*= 3 for each group, 3-5 IVDs were analyzed/mouse. Bars represent mean ± SD. *p≤0.05, Student’s *t* Test).

**Supplemental Figure 7:**
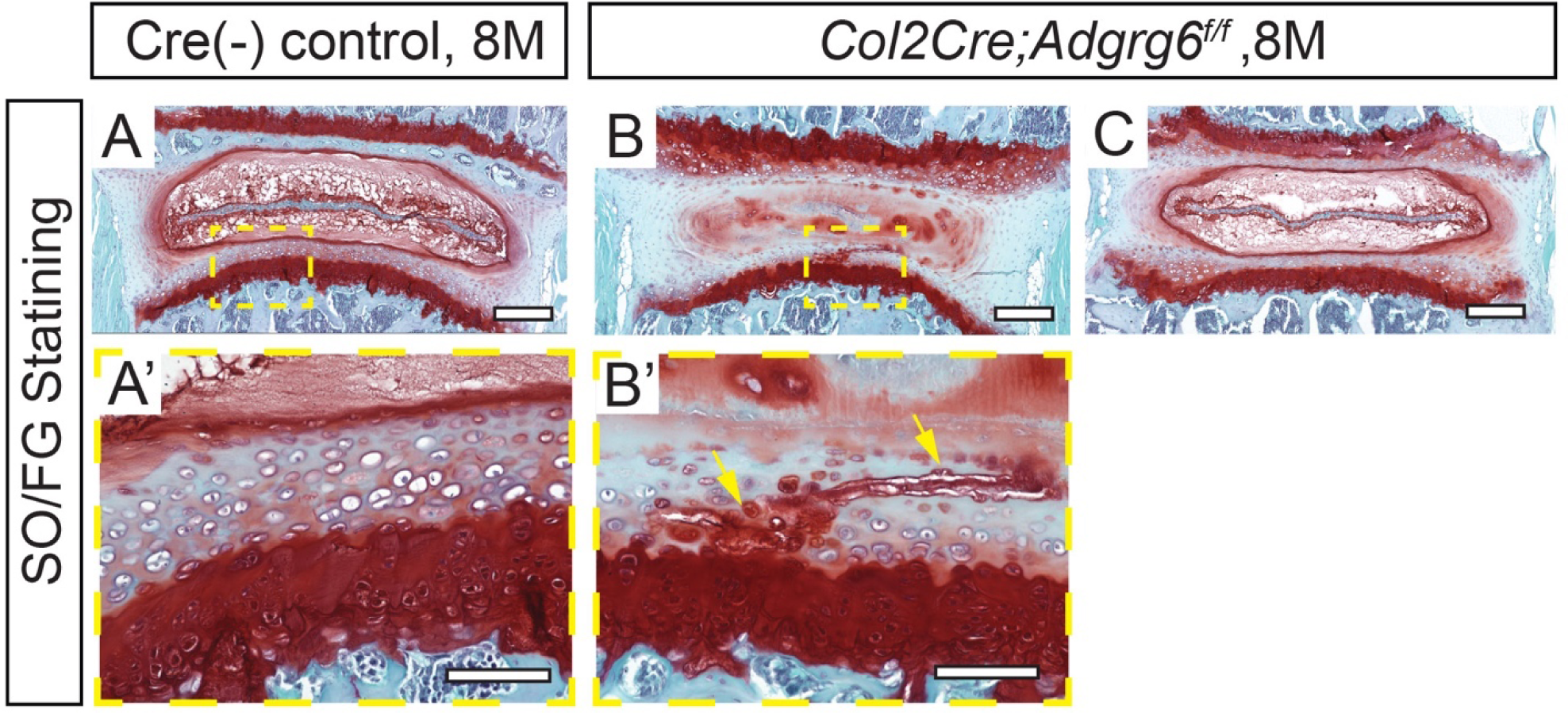
*Col2Cre;Adgrg6^f/f^* mutant mice display endplate-herniation of the IVD. (A-C) Representative medial-sectioned mouse IVDs stained with Safranin-O/Fast green (SO/FG) of Cre (-) control (A and A’) and *Col2Cre;Adgrg6^f/f^* mutant (B, B’, and C) mice by the age of 8 months. Endplate-oriented herniations is indicated with yellow arrows. These herniations are very hard to be captured by traditional two-dimensional histological analysis (B, which is out of the typical plane of section). C is an earlier section of the same mutant IVD as shown in B, looking completely normal. (*n*=3 for each group.) Scale bars: 200μm in (A-C), and 50μm in (A’, B’).

**Supplemental Figure 8:**
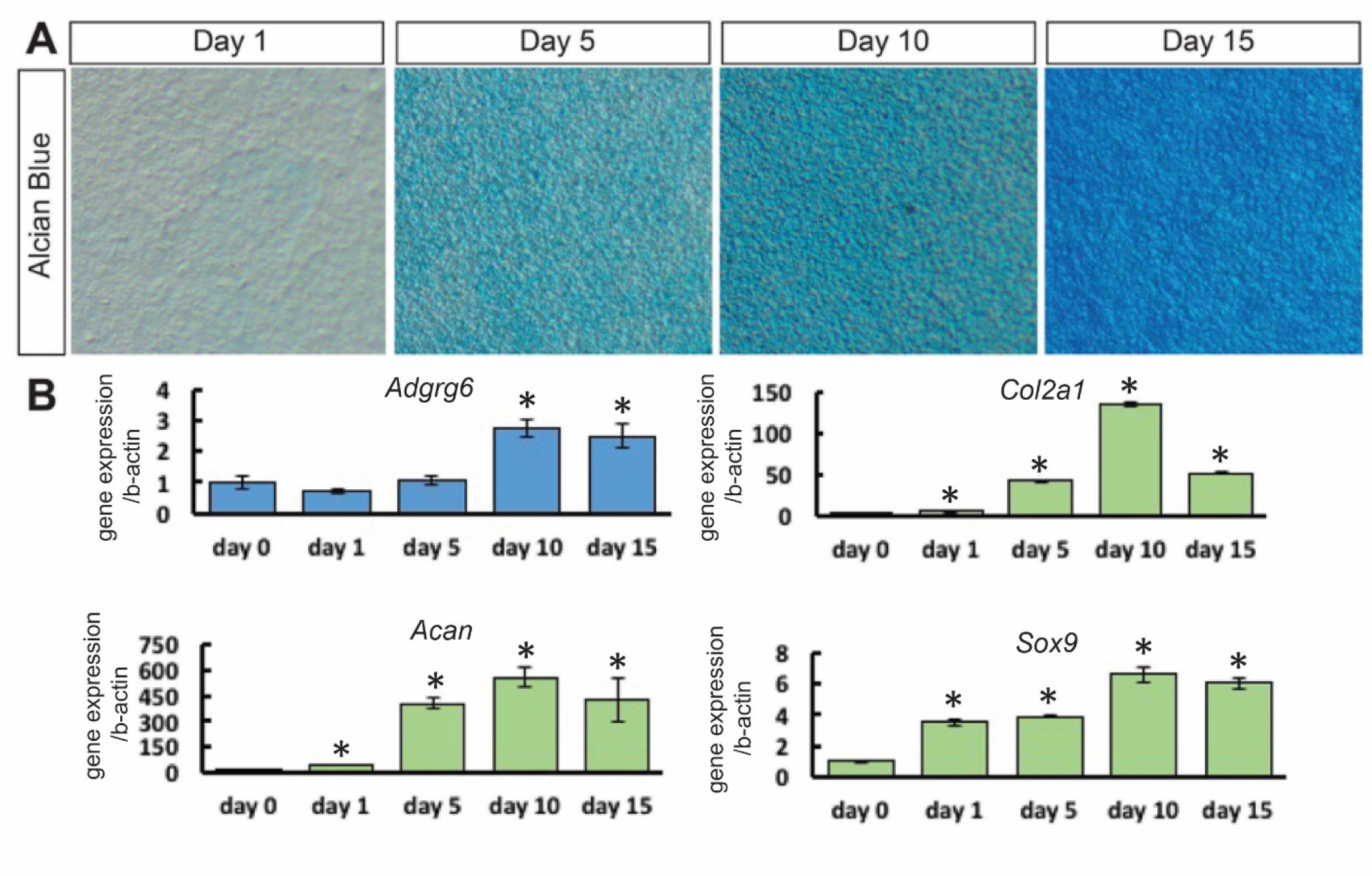
*Adgrg6* regulates ATDC5 cell maturation. (A) Alcian blue staining on ATDC5 cell culture during the maturation process. (B) Expression profiles of *Adgrg6*, *Col2a1*, *Acan*, and *Sox9* during ATDC5 cell maturation. The expression level of *Adgrg6* was gradually increased alone with other chondrogenesis markers including *Col2a1*, *Acan*, and *Sox9*. (*n*= 3 biological replicates and representative result is shown. Bars represent mean ± SD. *p≤0.05, Student’s t Test).

**Supplemental Figure 9:**
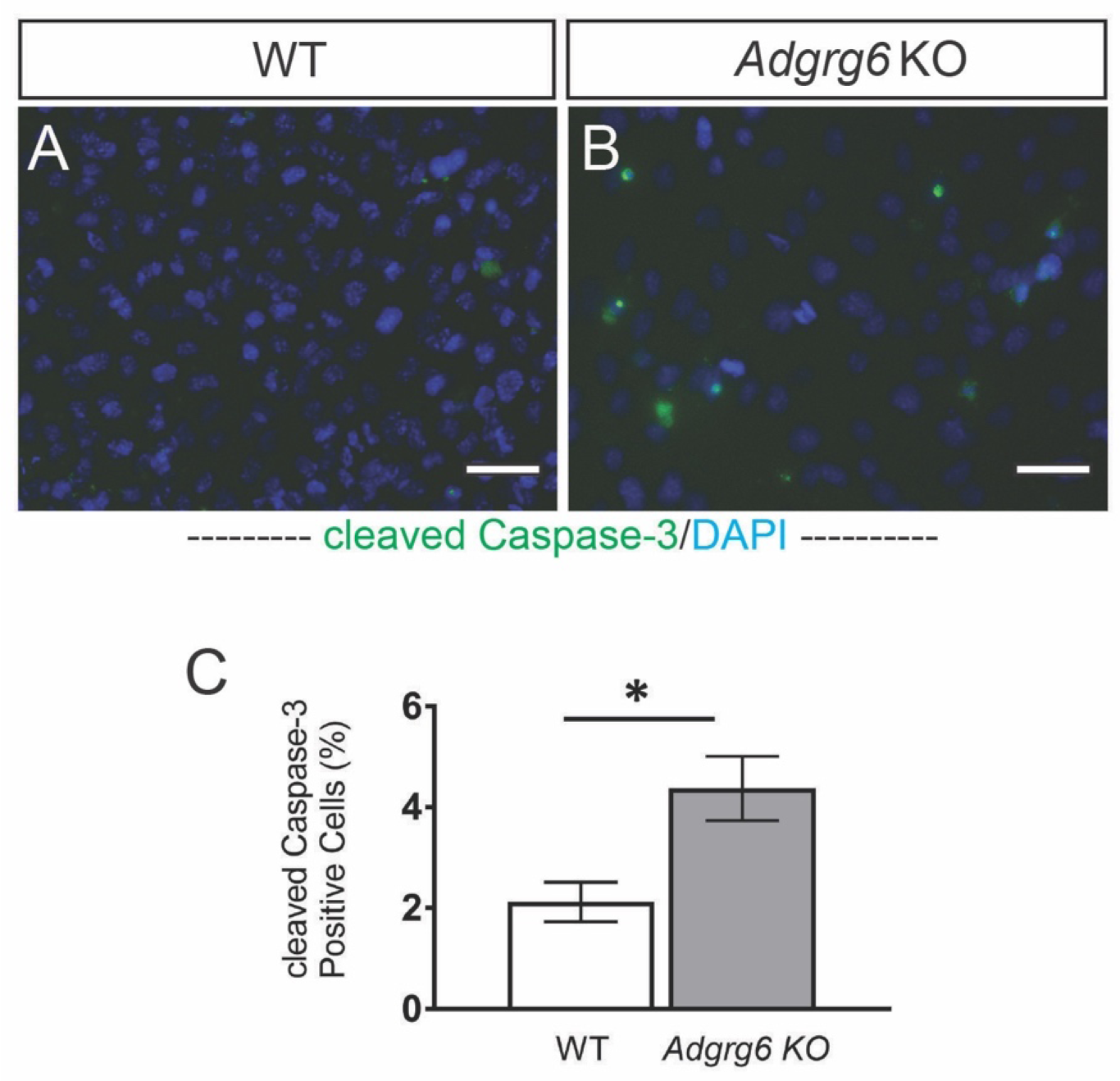
*Adgrg6* KO cells showed increased expression of apoptosis marker during maturation. (A, B) Immunofluorescence against cleaved-Caspace-3 (green) and DAPI staining (blue) and (C) quantification showing increased apoptosis in *Adgrg6* KO cells during maturation (10 days). (*n*= 3 biological replicates and representative result is shown. Bars represent mean ± SD. *p≤0.05, Student’s t Test). Scale bars: 50μm in (F, G).

**Supplemental Figure 10:**
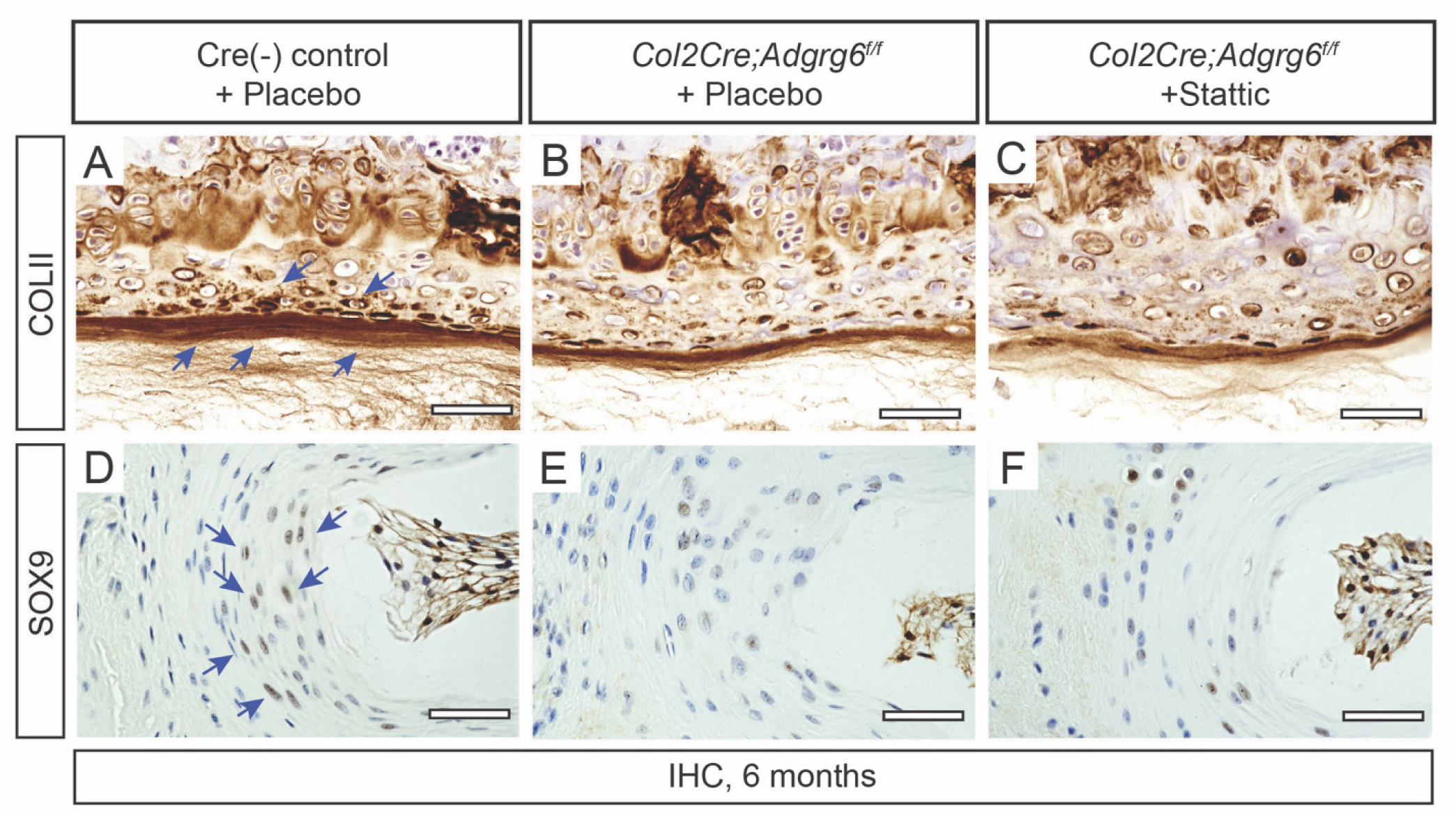
Inhibition of STAT3 activation by Stattic does not affect the expression of COLII and SOX9 in *Col2Cre;Adgrg6^f/f^* mutant mice. (A-F) IHC analysis of COLII and SOX9 in 6-month-old Cre (-) control (A, D) and *Col2Cre;Adgrg6^f/f^* mutant mice with (C, F) or without (B, E) Stattic treatment. Reduced COLII and SOX9 expression was observed in *Col2Cre;Adgrg6^f/f^* conditional mutant mice compared with Cre (-) control (A, D, blue arrows), but no significant improvement was observed after Stattic treatment (C, F). (*n*=3 for each group). Scale bars: 50μm in (A-F).

## Supplemental Tables

Table S1. Differential gene expression RNA-sequencing analysis from P20 IVD derived from *Col2Cre;Adgrg6^f/f^* and Cre(-) wild-type littermates.

Table S2. GO term analysis of P20 IVD derived from *Col2Cre;Adgrg6^f/f^* and Cre(-) wild-type littermates.

**Table S3.**
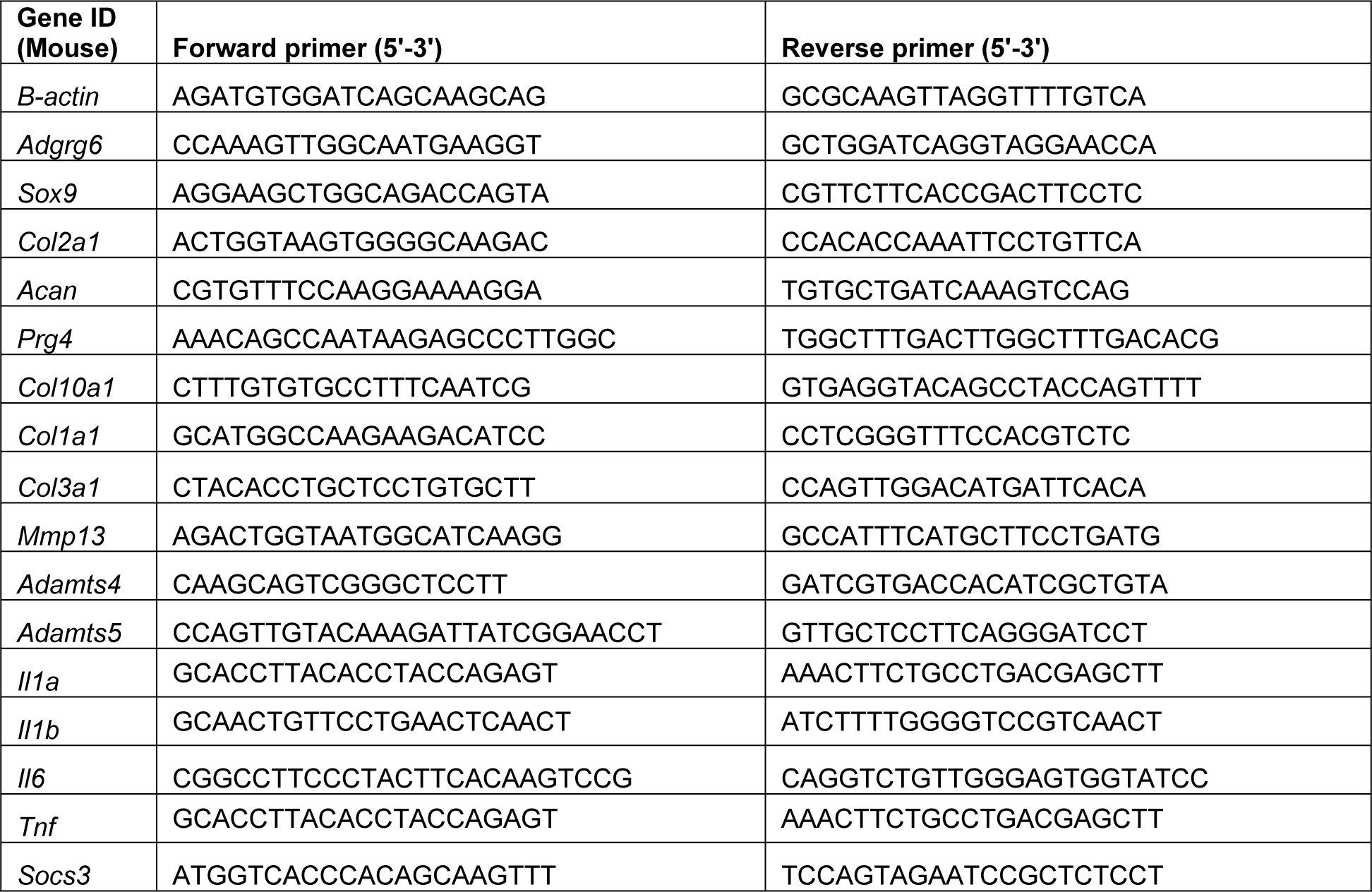
qPCR primers used in this study.

